# IGF targeting perturbs global replication through ribonucleotide reductase dysfunction

**DOI:** 10.1101/2020.08.11.245886

**Authors:** Guillaume Rieunier, Xiaoning Wu, Letitia Harris, Jack Mills, Ashwin Nandakumar, Laura Colling, Elena Seraia, Stephanie B. Hatch, Daniel Ebner, Lisa K. Folkes, Ulrike Weyer-Czernilofsky, Thomas Bogenrieder, Anderson J. Ryan, Valentine M. Macaulay

## Abstract

IGF receptor (IGF-1R) inhibition delays repair of radiation-induced DNA double-strand breaks (DSBs), prompting us to investigate whether IGF-1R influences endogenous DNA damage. We demonstrate that IGF-1R inhibition generates endogenous DNA lesions protected by 53BP1 bodies, indicating under-replicated DNA. We detect delayed replication fork progression in IGF-inhibited or IGF-1R depleted cancer cells, with activation of ATR-CHK1 signaling and the intra-S-phase checkpoint. This phenotype reflects unanticipated regulation of global replication by IGF-1, mediated via AKT, MEK/ERK and JUN to influence expression of ribonucleotide reductase (RNR) subunit RRM2. IGF-1R inhibition or depletion downregulate RRM2, compromising RNR function and dNTP supply. The resulting delay in fork progression and hallmarks of replication stress are rescued by RRM2 overexpression, confirming RRM2 as the critical factor through which IGF-1 regulates replication. Following targeted compound screens, we identify synergy between IGF inhibition and ATM loss, with evidence that IGF inhibition compromises growth of ATM null cells and xenografts. This synthetic lethal effect reflects conversion of single-stranded lesions in IGF-inhibited cells into toxic DSBs upon ATM inhibition. These data implicate IGF-1R in alleviating replication stress, and the identified reciprocal IGF:ATM co-dependence provides an approach to exploit this effect in ATM-deficient cancers.

## Introduction

Type 1 insulin-like growth factor receptor (IGF-1R) is a receptor tyrosine kinase (RTK) that is expressed by most common cancers, and promotes proliferation, cell survival and invasion (1). These effects are mediated via activation of multiple signalling pathways of which the most well-characterized are the phosphatidylinositol 3-kinase – AKT – mammalian target of rapamycin (PI3K-AKT-mTOR) and mitogen-activated protein kinase kinase – extracellular signal-regulated kinase (MEK-ERK) pathways (2-4). IGF-1R is required for cellular transformation by oncogenes including SV40 and RAS, and can itself induce transformation when overexpressed (5, 6).

We and others found that IGF-1R overexpression associates with clinical radioresistance in several types of cancer including breast and prostate cancers (7, 8). This appears to be a causative association in light of evidence from groups including ours that IGF-1R depletion or inhibition sensitizes to DNA damaging cytotoxic drugs and ionizing radiation (IR) *in vitro* and *in vivo* (9-15). We further reported that IGF-1R inhibition delays repair of IR-induced DNA double strand breaks (DSBs) and inhibits homologous recombination (HR) and non-homologous end-joining (NHEJ) in repair reporter assays (16). The precise molecular mechanisms linking the IGF axis to repair of exogenous DNA damage are not fully characterized, and appear independent of the well-characterized ability of IGFs to protect from apoptosis (16). Proteins and pathways that may be involved include IGF-regulated interaction between the IGF-1R docking protein insulin receptor substrate-1 (IRS-1) and RAD51 (17), and IGF-1R induced activation of the PI3K-AKT-mTOR and MEK-ERK pathways, both reported to mediate radioresistance (18-20).

There is clearly a substantial body of evidence supporting a role for the IGF axis and its downstream effectors in the response to DNA damage induced by IR. However, IGFs have not previously been implicated in maintaining replication integrity in the absence of exogenous DNA damage. We previously reported that IGF-1R inhibited or depleted tumor cells accumulate γH2AX foci, representing initial evidence of endogenous DNA lesions (21), but (22) the origins of these endogenous lesions were unknown. Foci formed by γH2AX indicate that atypical histone protein H2AX has undergone S139 phosphorylation at sites of damage by the PI3K-like kinases ataxia telangiectasia mutated (ATM), ataxia telangiectasia and Rad3-related protein (ATR), and DNA-dependent protein kinase (DNA-PK) (23, 24). DNA-PK and ATM are required for formation of γH2AX foci in response to IR-induced DSBs, ATM can be activated by diverse DNA structures in addition to DSBs, while ATR is required for response to extended single-stranded DNA (ssDNA) (23, 25, 26). Physiological processes that induce DSBs, which include V(d)J and meiotic recombination, do not occur in epithelial cancer cells, but extended ssDNA can be physiologically generated when DNA double helices separate to allow transcription or replication (27).

Here, we identify the source and functional consequences of endogenous DNA lesions in cells where IGF signalling is suppressed. We discover unanticipated regulation by IGF-1R of ribonucleotide reductase (RNR) function and dNTP supply; as a consequence, IGF axis blockade compromises completion of global DNA replication, delaying replication fork progression and allowing under-replicated DNA to progress through mitosis. We also reveal that replication stress in IGF-inhibited cells is maintained at a tolerable level by the important role of ATM in preventing conversion of single strand breaks (SSBs) into DSBs. Thus, replication stress in IGF-inhibited cells associates with a state of reciprocal co-dependence on ATM that can be exploited in therapy.

## Results

### IGF axis inhibition induces endogenous DNA damage and replication stress

To explore the role of IGF-1R in protection from endogenous DNA damage we used *IGF1R* gene silencing and IGF neutralizing antibody xentuzumab (28). In initial experiments, we detected increased γH2AX foci in xentuzumab-treated MCF7 breast cancer cells, at xentuzumab concentrations that inhibited IGF-1R and AKT-S6 phosphorylation (Figure 1A-B). ERK phosphorylation was not inhibited in MCF7 cells although was reduced in other breast cancer cell lines (Supplementary Figure S1A). Xentuzumab also caused concentration-dependent increase in lesions marked by 53BP1 (Figure 1C), and cyclin A – 53BP1 co-staining indicated that both xentuzumab and IGF-1R depletion increased 53BP1 signal in G1 (cyclin A-negative) cells (Figure 1D). These findings suggest appearance in G1 of 53BP1 nuclear bodies, which form to protect incompletely-replicated DNA from erosion during mitosis (29).

**Figure 1.**
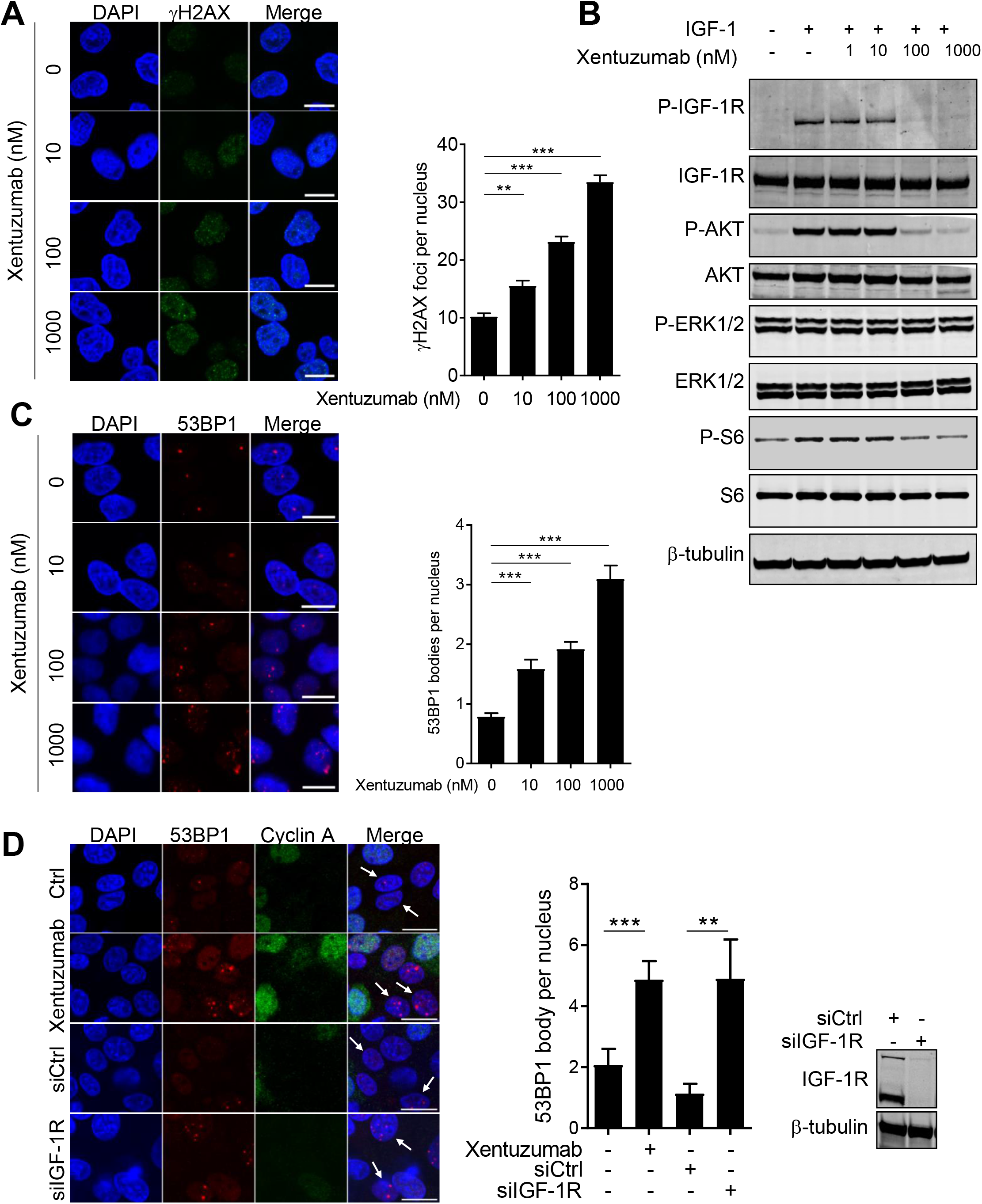
IGF-1R inhibition induces endogenous DNA damage. **A, C.** MCF7 cells were xentuzumab-treated for 5 days, showing representative immunofluorescence for: A, γH2AX; C, 53BP1 (scale bar 10μm). Graphs: mean ± SEM foci (n≥50 nuclei). **B.** Serum-starved MCF7 cells were xentuzumab-treated as A and with 50nM IGF-1 in final 15min. **D.** Cyclin A and 53BP1 immunofluorescence in MCF7 cells xentuzumab-treated (100nM) as A, or 72h post-siRNA transfection (scale 10μm, white arrows: cyclin A negative cells). Graph: mean ± SEM 53BP1 bodies, western blot confirms IGF-1R depletion.

The accumulation of 53BP1 bodies suggests a state of replication stress (30). This finding prompted us to perform DNA fiber assays, which revealed significantly shortened DNA tracts in IGF-1R-depleted MCF7 (Figure 2A-B) and KPL1 breast cancer cells (Supplementary Figure S1B). Replication fork speed was also reduced by IGF-1R depletion (Figure 2C), and comparable reduction in tract length and fork speed were detected within 6h exposure to xentuzumab (Figure 2D-E), suggesting a direct effect of IGF blockade. There was no increase in stalled or collapsed forks, identified as CldU (red) only signal or CIdU tracts followed by short IdU (green) tracts (Supplementary Fig S1C). To begin to investigate the cause of replication fork delay, we measured sister fork tracts, which extend in opposite directions from a unique replication origin. Sister forks progress at similar rates under physiological conditions or during replication stress that is not due to a physical impediment, and at different rates if one fork encounters DNA lesions (30, 31). We found no significant sister fork asymmetry upon IGF-1R depletion (Figure 2F), suggesting that replication stress originated from global reduction in replication, not obstruction by DNA lesions.

**Figure 2.**
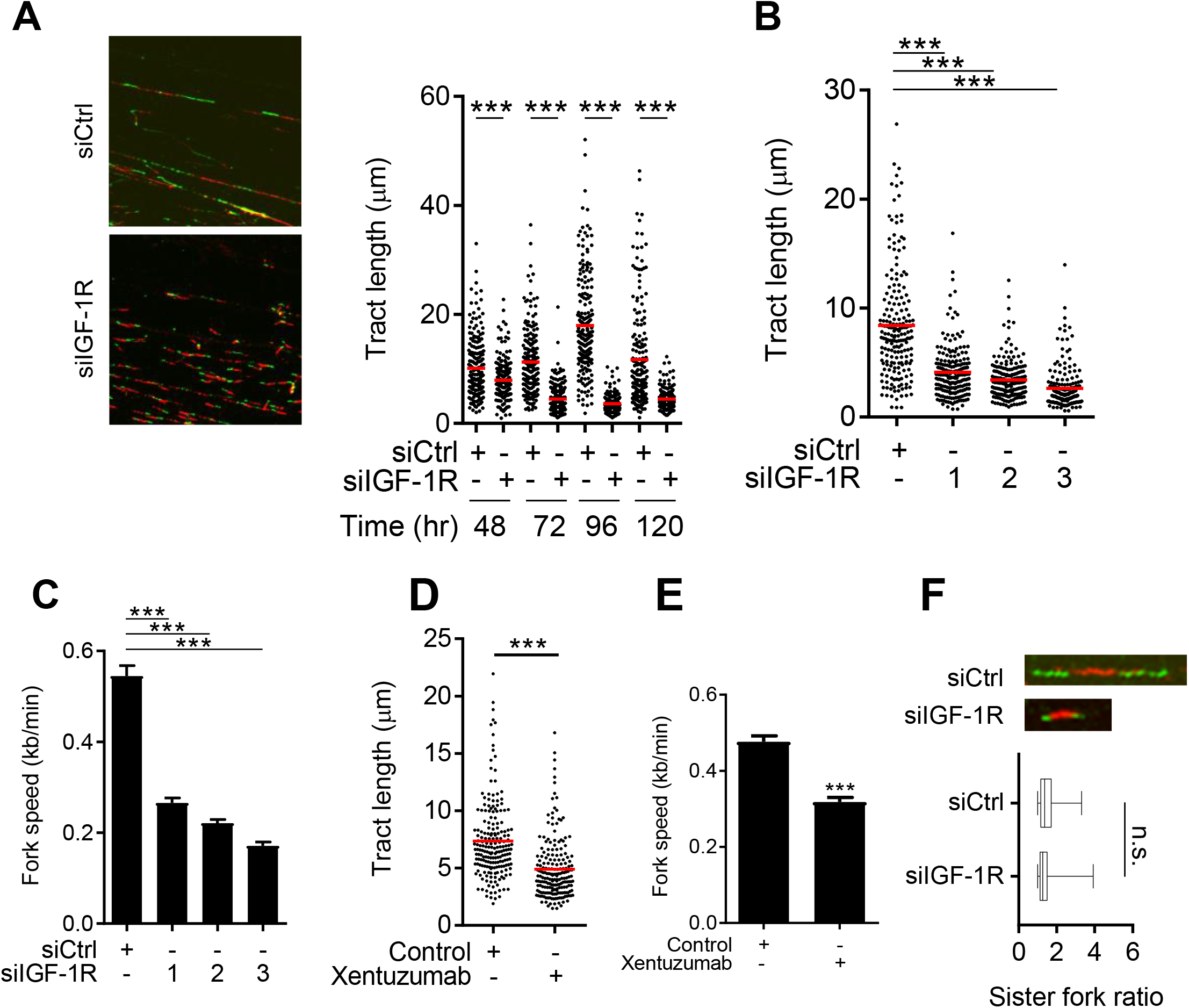
IGF axis targeting induces symmetrical reduction in replication fork progression. **A.** Replication tract images in MCF7 cells 72h after siRNA transfection. Graph to right: tract length (CIdU+IdU) in MCF7 cells 48-120h after siRNA-transfection, re-transfecting 96h and 120h time-points at 72h. Red line: median. **B, C.** MCF7 cells transfected with Control (siCtrl) or 3 different IGF-1R siRNAs, processed after 72h for DNA fiber analysis, showing tract length (B), fork speed (G). **D, E.** DNA fiber analysis in MCF7 cells treated with PBS (control) or 1 μM xentuzumab for 6h, showing tract length (H), fork speed (I). **F.** DNA fiber analysis comparing sister fork ratios. Upper panel: representative sister forks from MCF7 cells 72h post-siRNA transfection. Lower: sister fork ratios (median, lower/upper quartiles, minimum/maximum values).

The results of DNA fiber assays suggested limitation of factors required to complete replication. If this was indeed so, we predicted that regions of unreplicated single-stranded DNA (ssDNA) would trigger activation of the checkpoint kinase ATR (32). Increased phosphorylation on S33 of replication protein A (RPA) in IGF-1R depleted cells supported the presence of ssDNA (Supplementary Figure S1F). We next assessed S345-CHK1 phosphorylation, a marker of ATR activation at the replication fork, finding enhanced CHK1 phosphorylation in IGF-1R-depleted or xentuzumab-treated MCF7 and KPL1 cells (Figure 3A, Supplementary Figure S1D-E). Activation of ATR should lead to Inhibition of origin firing (33), and indeed newly-fired origins were reduced after IGF-1R depletion (Figure 3B). IGF-1R depletion also induced accumulation of cells in S-phase accumulation, detectable in asynchronous cultures (Supplementary Figure S1G) and exacerbated upon cell cycle synchronization (Figure 3C-D; Supplementary Figure S1H), representing further evidence of ATR activation. IGFs are well-known to promote progression through G1/S and G2/M checkpoints via cyclin upregulation, CDK activation and Rb phosphorylation (1). Our data show that by regulating fork progression, IGFs also influence the rate of transition through S-phase, and reveal hallmarks of replication stress when IGF-1R is non-functional.

**Figure 3.**
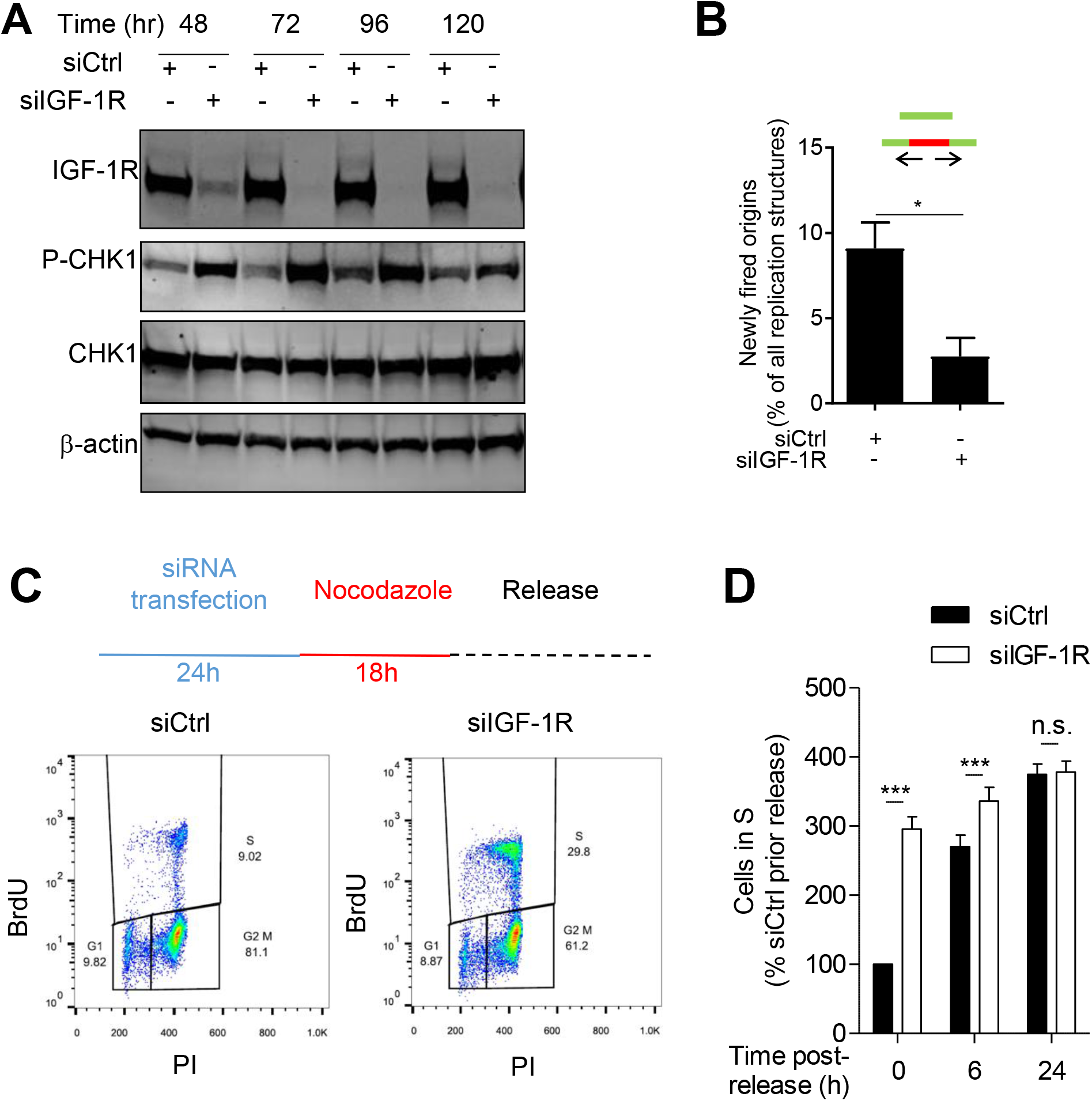
IGF-1R depletion activates ATR-CHK1 and intra-S checkpoint. **A.** MCF7 cells were siRNA transfected and after 72h were processed for DNA fiber analysis to quantify newly-fired origins. Upper panel: two types of newly-fired origins (arrows, direction of fork progression). **B.** MCF7 cells were siRNA transfected as A), harvested after 48-120h for western blot for phospho-S345 and total CHK1. Cells were re-transfected at 72h. **C-D.** MCF7 cells were siRNA-transfected, the following day treated with nocodazole, after 18h released into nocodazole-free medium and collected after 0-24h for cell cycle analysis (n=3 independent experiments).

### IGF-1 regulates RRM2 expression and RNR activity

Having identified replication fork delay and shortened but symmetrical sister forks in IGF-1R depleted cells (Figure 2A-B, F), we hypothesized that IGFs regulate the supply of factors required to complete replication. Quantification of deoxyribonucleoside triphosphates (dNTPs) revealed significant reduction in dATP content in IGF-1R-depleted MCF7 cells, while dTTP and dGTP were unchanged and dCTP undetectable (Figure 4A; Supplementary Figure S2A-B). All dNTPs were detected in HCT15 colorectal cancer (CRC) cells, which proliferate more rapidly than MCF7; dATP content was reduced by IGF-1R depletion as in MCF7, while dCTP, dGTP and dTTP were unaltered (Figure 4B). These differential changes led us to investigate effects on RNR, which converts ribonucleotide diphosphates (NDPs) to deoxyribonucleotides (dNDPs), the rate limiting step for dNTP production (34). We noted that a microarray study in MCF7 cells had generated an IGF gene signature that associates with adverse outcome in breast cancer, and included upregulation of cyclins, components of the replication machinery and the regulatory subunit M2 (RRM2) of RNR (35). In the same cell line, we found that RRM2 mRNA and protein were downregulated by IGF-1R depletion and xentuzumab, while catalytic subunit RRM1 was unaffected (Figure 4C-E; Supplementary Figure S2C). RRM2 was also downregulated in IGF-1R depleted KPL1, HCC1143, ZR-75-1 and T47D breast cancer, DU145 prostate cancer and U2OS osteosarcoma cells (Supplementary Figure S2D). Consistent with accumulation of S phase cells (Figure 3D), cyclin A was upregulated by IGF-1R depletion (Figure 4C). Consistent with the report of Lee and colleagues (35), *RRM2* was upregulated at the mRNA level by IGF-1 (Figure 4F), and to investigate the regulators of RRM2 expression we inhibited the principal effectors downstream of IGF-1R. The activities of MEK inhibitor trametinib and AKT inhibitor AZD5363 were confirmed by reduction in ERK and S6 phosphorylation respectively, and both agents were found to suppress IGF-induced RRM2 upregulation (Figure 4G-H). We and others reported that nuclear IGF-1R undergoes nuclear translocation and recruitment to chromatin (36-38), with evidence from our ChIP-seq of IGF-1R on proximal promoters (39). However, we did not detect IGF-1R at the *RRM2* promoter (Supplementary Figure S2E), suggesting that IGFs regulate RRM2 via canonical signalling, not via non-canonical nuclear translocation. The RRM2 promoter contains binding motifs for multiple transcription factors (www.genecards.org/cgi-bin/carddisp.pl?gene=RRM2), including E2F and the FOS/JUN dimer AP-1. To investigate the contribution of these candidate regulators of RRM2 expression downstream of MEK-ERK, we knocked down E2F and JUN. In E2F-depleted MCF7 cells there was evidence of RRM2 downregulation, and co-depletion of both IGF-1R and E2F achieved more profound RRM2 downregulation than was achieved by depleting either factor alone (Supplementary Figure S2F). This suggests that IGF-1R was not functioning in the same pathway as E2F with respect to RRM2 regulation. JUN depletion also downregulated RRM2, comparable to the degree of RRM2 downregulation induced by depleting IGF-1R, and here the effect of IGF-1R:JUN co-depletion achieved no greater RRM2 downregulation than effects of knocking down IGF-1R or JUN separately (Figure 4I). This result suggested JUN as a candidate mediator of IGF effects on RRM2 expression. This finding is consistent with the presence of JUN consensus binding motif TGACTCA (40) within the sequence of the *RRM2* promoter, and with the known function of JUN as a downstream transcriptional effector of MEK-ERK signalling (41) To further explore the role of JUN in regulating *RRM2* promoter activity we transfected MCF7 cells with a luciferase reporter incorporating the *RRM2* promoter, and performed depletion epistasis analysis using luciferase activity as a readout. *RRM2* promoter activity was reduced by depletion of either IGF-1R or JUN, and as with the assessment of protein levels (Figure 4I), promoter activity was not further reduced by depleting both factors (Figure 4J). These analyses suggest an epistatic relationship between IGF-1R and JUN, and support a model in which IGFs regulate RRM2 expression at least in part via MEK-ERK-JUN signalling.

**Figure 4.**
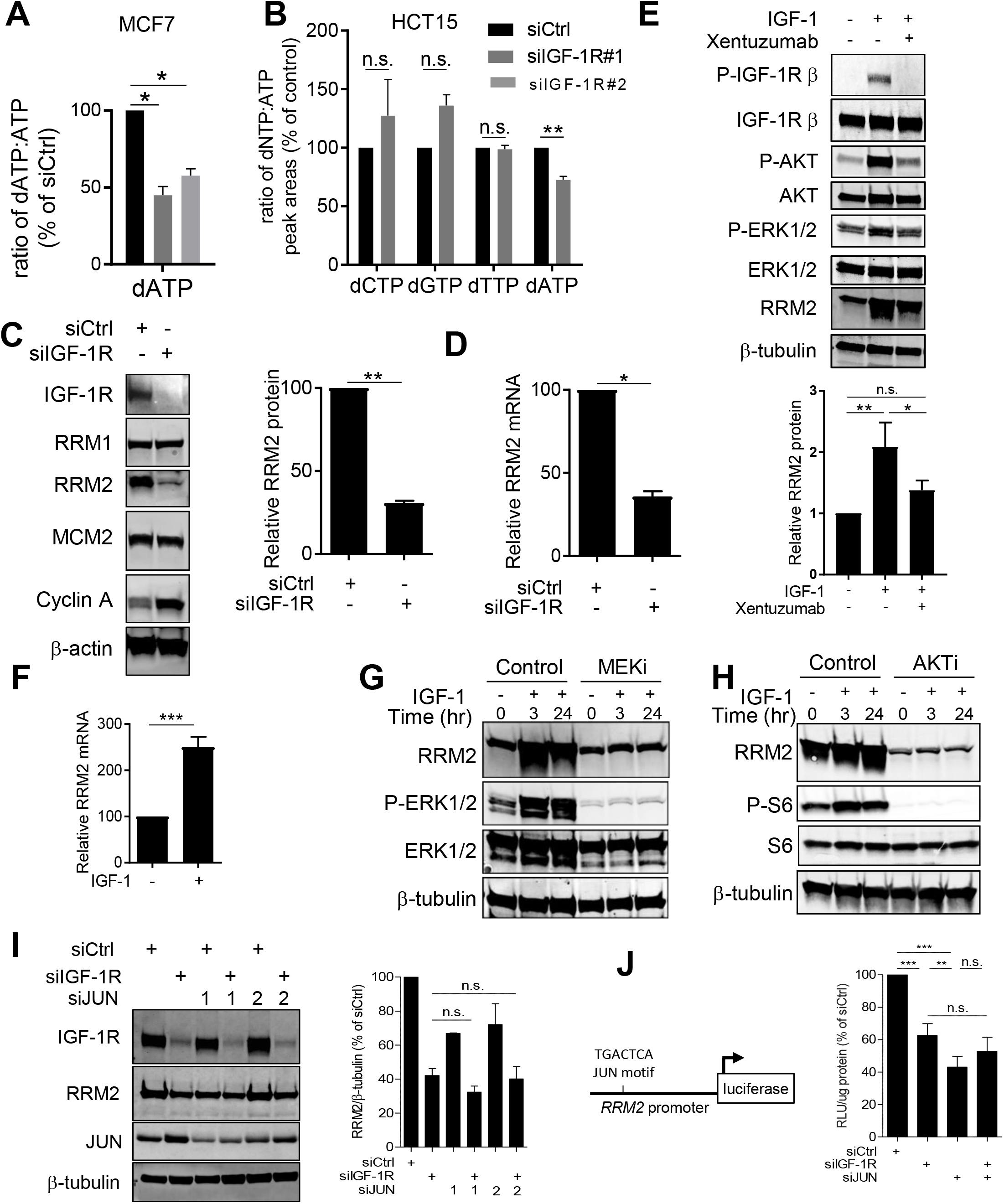
IGF-1R regulates RNR activity via AKT, MEK-ERK and JUN. **A, B.** Mean ± SEM fold-change in dNTP content in A, MCF7; B, HCT15 cells 48h after siRNA-transfection. **C, D.** MCF7 cells were siRNA transfected, collected after 48h for: C, western blot, graph shows mean ± SEM RRM2 protein (n=3 western blots); D, qPCR for *RRM2* mRNA. **E.** Serum-starved MCF7 cells pre-treated with 1μM xentuzumab for 2h then xentuzumab with 50nM IGF-1 for 24h. Graph below: mean ± SEM RRM2 protein (n=3 blots). **F.** Serum-starved MCF7 cells IGF-treated for 24h, *RRM2* mRNA assessed by qPCR. **G, H.** Serum-starved MCF7 cells treated with IGF-1 alone or with 10nM MEK inhibitor (MEKi) trametinib (G) or 3.5mM AKT inhibitor (AKTi) AZD5363 (H). **I-J.** MCF7 cells transfected with siIGF-1R and/or siJUN were analysed by: I, western blot; J: luciferase assay after transient transfection with *RRM2* promoter luciferase reporter (n=3).

We used two models to investigate the importance of RRM2 regulation to the replication stress phenotype induced by IGF-1R depletion. Firstly, we utilized RRM2 stably-overexpressing U2OS cells that had been generated by D’Angiolella and colleagues (42), first confirming constitutive RRM2 overexpression that was resistant to IGF-1R knockdown (Figure 5A). In control (empty vector) transfectants, IGF neutralizing antibody xentuzumab induced phosphorylation of ATR targets CHK1 and RPA (Figure 5B), consistent with results in MCF7 (Supplementary Figure S1D-E), while RRM2 overexpression suppressed CHK1 and RPA phosphorylation. We then quantified 53BP1 bodies as surrogate markers of under-replicated DNA, finding increased 53BP1 bodies in IGF-1R depleted U2OS control cells, again consistent with data in MCF7 (Figure 1D). However, this increase upon IGF-1R depletion was completely prevented by RRM2-overexpression (Figure 5C). We also assessed the role of RRM2 in a second model, stably-transfecting MCF7 cells with control vector or RRM2 cDNA, and used these cells to perform DNA fiber assays and cell cycle analysis. In control transfectants, IGF-1R depletion caused significant shortening of DNA tracts and modest but statistically significant accumulation of cells in S-phase (Figure 5E, Supplementary Figure S2G) comparable to effects in parental MCF7 cells (Figure 2A-B, Figure 3C-D). In MCF7 cells stably over-expressing RRM2 (Figure 5D) there was partial rescue from DNA tract shortening after IGF-1R depletion (Figure 5E) and no accumulation of cells in S phase (Supplementary Figure S2G). Rescue from the hallmarks of replication stress is consistent with data on 53BP1 bodies in the U2OS model (Figure 5C), with partial rescue in MCF7 likely attributable to the reduction in endogenous RRM2 upon IGF-1R depletion (Figure 5D). This rescue identifies RRM2 as a central factor in the replication stress phenotype induced by IGF-1R depletion.

**Figure 5.**
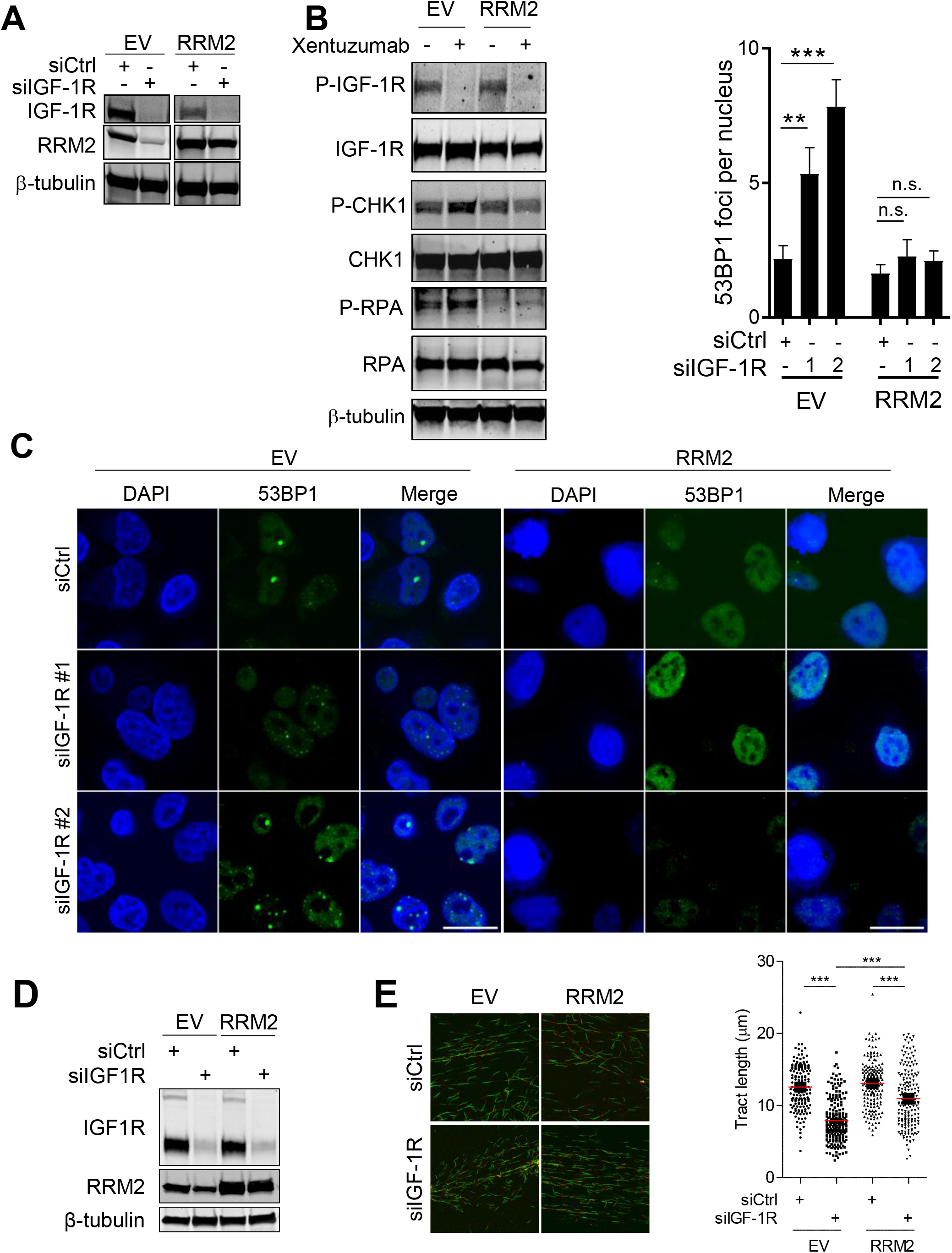
RRM2 overexpression rescues IGF-1R depleted or inhibited cells from replication stress. **A-B.** U2OS cells stably-transfected with RRM2 or empty vector (EV) were analysed by western blot 48h after IGF-1R siRNA transfection (A) or 72h treatment with 100nM xentuzumab (B). **C.** Cells transfected as A were processed for 53BP1 immunofluorescence (scale 10μm). Graph above: mean ± SEM 53BP1 bodies (n=30 nuclei). **D.** Western blot of MCF7 cells stably transfected with empty vector (EV) or RRM2 cDNA, harvested 48h after siRNA transfection. **E.** Control and RRM2-transfected MCF7 cells were processed for DNA fiber analysis 48h after siRNA transfection showing: left, representative images; right, quantification of tract length (n=200).

### ATM loss is synthetically lethal to IGF-inhibited cells and tumors

It was clear that IGF-1R depletion or inhibition caused significant perturbation of replication fork progression, and we next quantified the associated effects on proliferation, with a view to assessing potential clinical relevance. Therefore, we tested xentuzumab in the five luminal breast cancer cell lines in which RRM2 was downregulated by IGF-1R depletion (Supplementary Figure S2D). We treated cells with xentuzumab at 1 μM, which approximates to the steady-state plasma concentration at the 1000 mg/week dose selected for Phase II evaluation (43). At this concentration, xentuzumab caused relatively little inhibition of cell viability (Figure 6A). This minor effect in the face of significant replication stress suggested existence of a backup pathway that alleviates toxic effects of replication intermediates, allowing a threshold of tolerable replication stress. To identify components of such a pathway, we performed a targeted compound screen using kinase, cell cycle, DNA replication and repair inhibitors in the five luminal breast cancer cell lines. We discuss MCF7 data here, and will report other outcomes separately (Wu, Rieunier et al in preparation). Assessment of the 12 top-ranked compounds achieving greatest viability inhibition with xentuzumab revealed partial overlap at the three screened compound concentrations (Figure 6B). Included here were inhibitors of PARP, EGFR and MEK (Supplementary Table S2), which were already identified in previous work by our group and others as showing potential for combination with IGF-1R blockade (22, 44-47). Of five compounds ranked highly at all concentrations (Supplementary Figure S3A), only one, ATM inhibitor KU-55933, had not previously been identified as an attractive partner for IGF co-inhibition. We validated KU-55933 as a screen hit in MCF7 and ZR-75-1 cells, finding that KU-55933 had little effect in control-treated cells, consistent with the known lack of toxicity of ATM inhibition in the absence of exogenous DNA damage (48), while addition of xentuzumab significantly inhibited viability at KU-55933 concentrations ≥300 nM (Figure 6C; Supplementary Figure S3B). We confirmed that at these concentrations, KU-55933 inhibited irradiation-induced phosphorylation of ATM target KAP1, both in the absence and presence of xentuzumab (Figure 6C-D; Supplementary Figure S3B-C). Xentuzumab also caused synergistic inhibition of the viability of MCF7, MiaPaca-2 and BXPC-3 cells when combined with a second ATM inhibitor, AZD0156 (Figure 6E, Supplementary Figure S3D-E) at clinically-achieved sub-micromolar concentrations (49). Furthermore, MCF7 cell death was significantly increased in cells exposed to xentuzumab with KU-55933 or AZD0156 (Figure 6G-H). Given evidence of ATR activation upon IGF-1R targeting (Figure 3A, Supplementary Figure S1D-F), we also tested for synthetic lethality with ATR inhibitors AZ20 or VE-821. However, we found no increased toxicity on addition of xentuzumab (Supplementary Figure S3F-G), supporting ATM-specific effects of the combination with KU-55933 or AZD0156.

**Figure 6.**
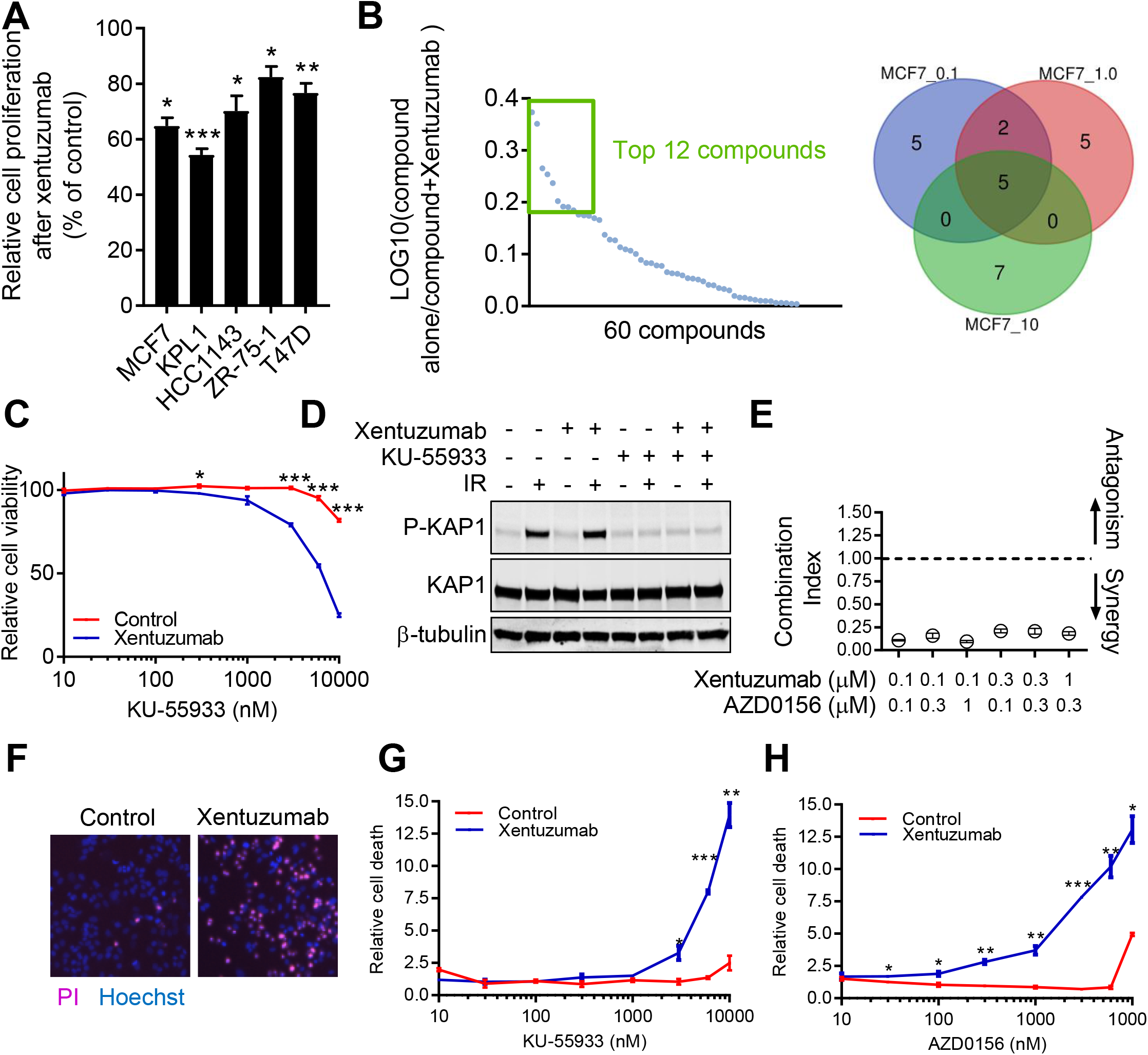
ATM maintains viability in the context of xentuzumab-induced replication stress. **A.** Relative viability after 5 days treatment with 1μM xentuzumab. **B.** MCF7 cells treated with 1 μM xentuzumab alone or with 60 compounds at 0.1, 1 and 10 μM, viability measured after 5 days. Correlation between duplicate screens was 0.97, and screen Z-Factors calculated using positive control for viability inhibition (PLK1 inhibitor BI-2536) were 0.64 at 100 nM, 0.68 at 1 μM and 0.72 at 10 μM, indicating high-quality screens. Graph: compounds screened at 0.1 mM, ranked by ratio of log10 compound alone/ compound+xentuzumab. Green box: top 12 compounds inducing at least additive effects (log10 ratio>0). Venn diagram: top-ranked hits in common at 0.1, 1 and 10 μM compound (also see Supplemental Figure S4A). **C.** MCF7 viability treated for 5 days with KU-55933 alone/with 1μM xentuzumab. **D.** MCF7 cells pre-treated with 10μM KU-55933 and/or xentuzumab for 72h, analysed 30 min after 5Gy irradiation. **E.** Combination indices (CI) calculated from viability data. CI values <0.8 indicate synergy and <0.3, strong synergy. **F-H.** Cell death measured in MCF7 cells as PI-positive cells as % of total (Hoechst-positive) cells, expressed relative to solvent-treated controls, showing F: representative appearances, **G-H.** relative cell death after treatment of MCF7 cells for 5 days with 1μM xentuzumab and KU-55933 or AZD0156.

*ATM* mutations and deletions are detectable in a significant proportion of epithelial cancers including CRC (12%), lung (10%), prostate (7%), pancreatic (4%) and breast (3.5%) cancers ((50, 51), cbioportal.org). Furthermore, loss of ATM protein is reported in up to 60% of sporadic breast cancers and ~13% of pancreatic ductal adenocarcinomas (PDAC) (52, 53), significantly exceeding the rate of *ATM* mutation, reflecting complex regulation of ATM expression. Given this evidence of frequent ATM loss, we took four approaches to test IGF dependence in ATM deficient cells. First, we accessed publicly-available data from 926 cancer cell lines (cancerrxgene.org, kobic.kr), including cell lines harboring homozygous *ATM* mutations, which lack functional ATM. Homozygous *ATM*^−/−^ cell lines were more sensitive to IGF-1R inhibitor BMS-754807 (p=0.0018), with a trend to increased sensitivity to linsitinib (p=0.08, Figure 7A, Supplementary Figure S4A).

**Figure 7.**
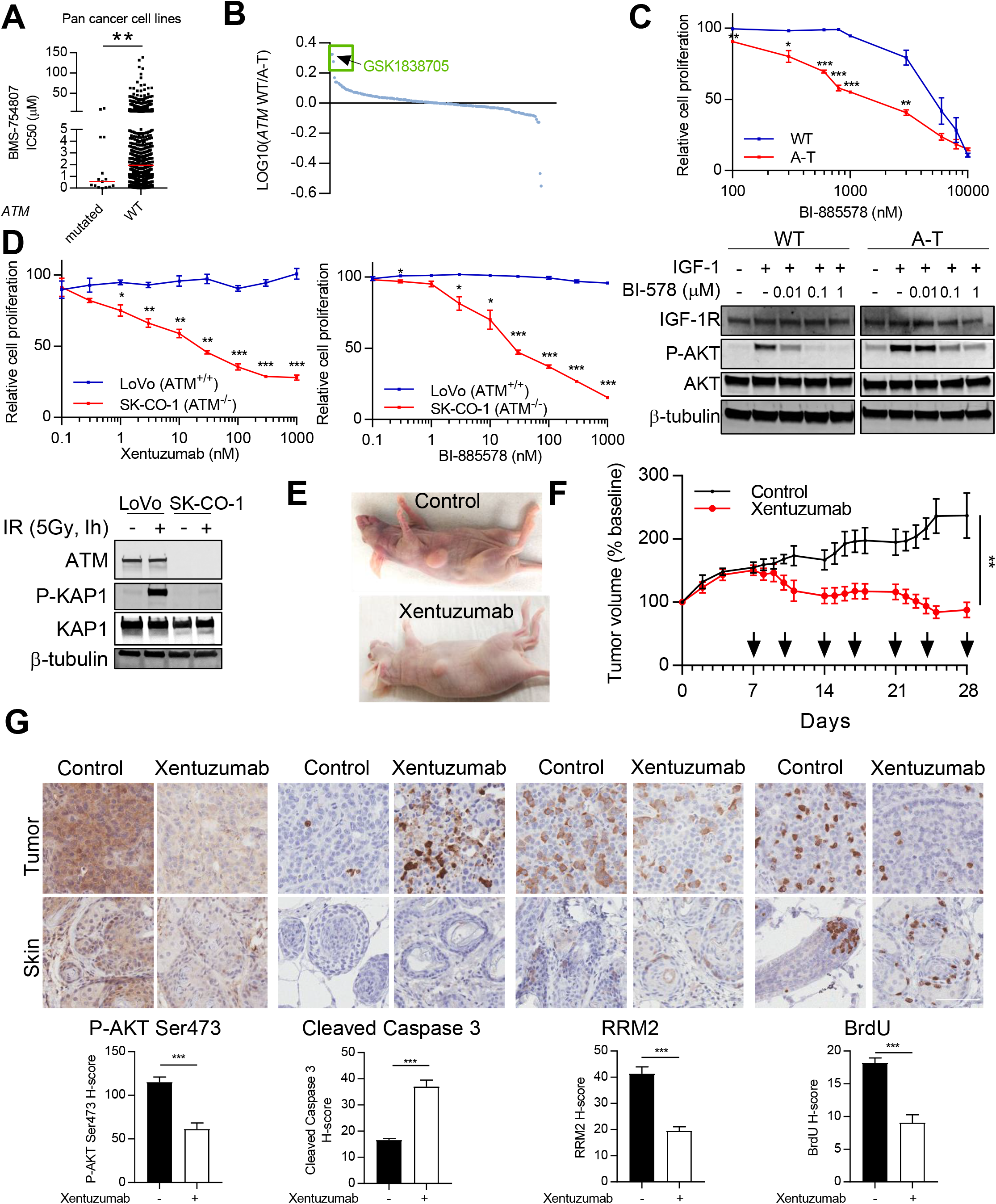
ATM loss sensitizes to IGF inhibition. **A.** Pan-cancer analysis (926 cell lines) associating *ATM* homozygous mutations and sensitivity to IGF-1R inhibitor BMS-754807 (cancerxgene.org). **B.** *ATM*^+/+^ and *ATM*^−/−^ fibroblasts treated with 188 kinase inhibitors (1μM), viability measured after 3 days, compounds ranked by ratio of log10 viability. Green box: top-ranked compounds, top hit arrowed. **C.** Fibroblasts as B treated with BI-885578, viability measured after 5 days. Blot below: serum-starved fibroblasts treated with BI-885578 for 24h and in final 15 min with IGF-1. **D.** Viability of CRC cells treated with xentuzumab or BI-885578 for 5 days. Blot below confirms ATM null status of SK-CO-1. **E-F.** Mice bearing SK-CO-1 xenografts treated twice weekly (arrows) with solvent or xentuzumab showing E: representative tumor-bearing mice, F: tumor volumes. **G.** Immunohistochemical analysis of SK-CO-1 tumors and mouse skin (scale 100μm). H-scores for RRM2 and BrdU represent moderate-strong (2-3+) staining.

Secondly, we took an unbiased approach to compare IGF-1R function in *ATM*^+/+^ and null (A-T) fibroblasts (Figure 7B). After confirming that *ATM*^−/−^ fibroblasts lacked functional ATM (Supplementary Figure S4B), we used these cells in a kinase inhibitor screen. Consistent with identification of ATM as a hit in the xentuzumab screen (Figure 6B-C), IGF-1R inhibitor GSK1838705 was the top synthetic lethal interaction in *ATM*^−/−^ fibroblasts when testing compounds at 1 μM, and the second-ranked hit at 0.1 μM (Supplementary Figure S4C). Low-throughput assays confirmed that *ATM*^−/−^ fibroblasts were significantly more sensitive to IGF-1R kinase inhibitor BI-885578 (54) and to xentuzumab (Figure 4C; Supplementary Figure S4D). BI-885578 suppressed IGF-induced AKT activation in both ATM proficient and deficient cells (Figure 7C, lower panel), so relative resistance of *ATM*^+/+^ cells to IGF-1R blockade was not due to failure of target inhibition. Thirdly, we used an isogenic ATM proficient/deficient H322 lung cancer model (55), finding that *ATM*-null cells were significantly more sensitive to BI-885578 (Figure S4E). H322 cells are not tumorigenic (55), so as a fourth approach we sought alternative *ATM*-null models for *in vivo* testing. Given the frequency of *ATM* mutation/deletion in CRC, we used CRC cell lines LoVo and SK-CO-1, which both harbor WT *PTEN* and β-catenin and mutant *KRAS* and *APC* (54) but differ in ATM status, SK-CO-1 being ATM null (56). LoVo cells were resistant to IGF-1R inhibition, consistent with data in (54), despite effective suppression of IGF-induced AKT phosphorylation (Supplementary Figure S4F), while SK-CO-1 cells were significantly more sensitive to xentuzumab and BI-885578 (Figure 7D), suggesting that *ATM* loss increases dependence on IGF-1R.

We next questioned whether IGF-1R influences growth of *ATM*-null tumors *in vivo*. Tumor cell proliferation is slower *in vivo* than *in vitro*, but it is questionable whether DNA replication is also slower, and if so whether this would affect replication stress induction by IGF blockade. Therefore, we treated mice bearing SK-CO-1 xenografts with xentuzumab at the dose and schedule used previously (28). Xentuzumab was well-tolerated and caused significant inhibition of tumor growth (Figure 7E-F, Supplementary Figure S4G). By immunohistochemistry we confirmed that SK-CO-1 xenografts were ATM-null, mouse epidermis acting as positive control for ATM detection (Supplementary Figure S4H). Xentuzumab blocked phospho-Ser473 AKT in both human tumor and mouse skin (Figure 7G), as expected given cross-reactivity with murine IGFs (28). Cleaved caspase 3 signal was significantly increased in xentuzumab-treated tumors, with no evidence of apoptosis in normal mouse skin (Figure 7G), suggesting that xentuzumab may be unlikely to cause toxicity to ATM-proficient normal tissues. We then assessed RRM2 expression, which is known to be cell cycle-regulated, increasing during S phase (34), likely explaining heterogeneous signal in tumor tissue and low expression in skin, except in hair follicles. RRM2 was significantly downregulated in xentuzumab-treated tumors, particularly when considering moderate (2+) or strong (3+) RRM2 signal (Figure 7G), consistent with *in vitro* findings (Figure 4E). We also assessed BrdU incorporation to measure DNA replication *in vivo*. We found heterogeneous incorporation with strong signal in tumor cells and hair follicles (Figure 7G), with significant reduction in moderate-strong BrdU signal after xentuzumab treatment, consistent with replication defects observed in IGF-1R depleted or inhibited cells *in vitro* (Figure 2A-E). There was less striking reduction in RRM2 and BrdU when including weak (1+) positivity (Supplementary Figure S4I-J), consistent with the S phase arrest induced in MCF7 cells by IGF-1R depletion (Figure – asynchronous cell cycle data). Taken together, these data suggest that tumors lacking ATM may be responsive to IGF blockade, and provide a rationale to investigate ATM status in selecting patients for IGF inhibitor trials.

### Loss of functional ATM converts replication stress-associated SSBs in IGF inhibited cells into DSBs

Next, we explored the role of ATM in alleviating toxic effects of replication stress in IGF inhibited cells. We had speculated that ATM might be activated in response to DNA lesions induced in IGF-inhibited cells, and this activation could be protective. However, when we had earlier assessed ATM activity to confirm the bioactivity of KU-99533, we found no evidence that xentuzumab treatment or IGF-1R depletion induced KAP1 and ATM phosphorylation, indicating that ATM had not been activated (Figure 6D; Supplementary Figure S3H). As a first approach to assessing the contribution of ATM in protecting replication forks we performed DNA fiber assays on CRC cell lines LoVo and SK-CO-1, which are matched for major mutations and differ in ATM status. Upon IGF-1R depletion we observed significant shortening of DNA tracts in SK-CO-1 cells (16.1 ± 4μm to 8.7 ± 3.7μm; Figure 8A), consistent with the effect we showed in MCF7 and KPL1 breast cancer cells (Figure 2A-B; Supplementary Figure S1B). The basal rate of replication fork progression was slower in LoVo cells, and in contrast to the major effect in SK-CO-1 cells there was a smaller change upon IGF-1R depletion (10 ± 2.9μm to 9 ± 2.6μm; Figure 8A), suggesting a protective role for ATM.

**Figure 8.**
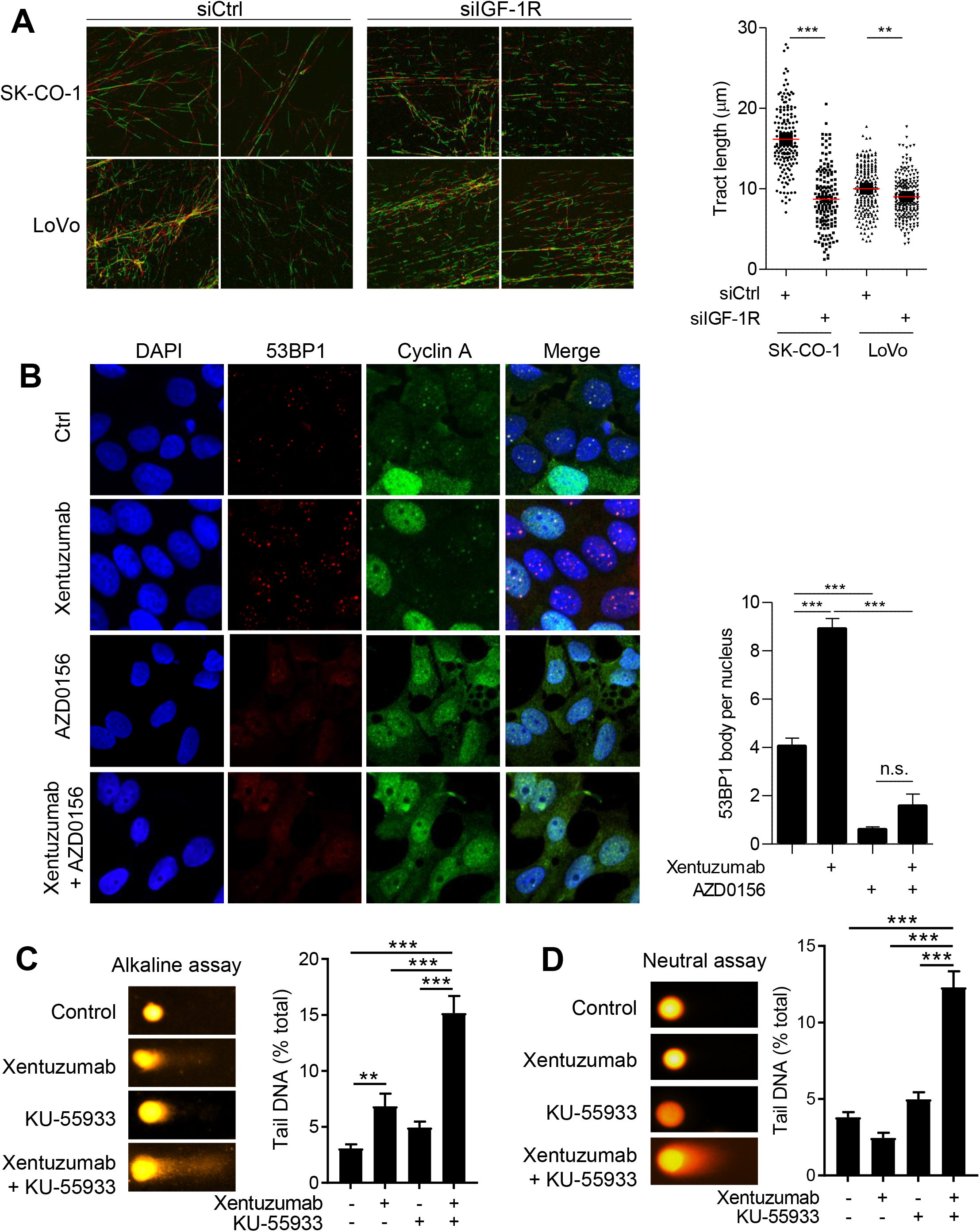
Tolerable ssDNA lesions in IGF-1R deficient cells are converted to DSBs by ATM loss. **A.** CRC cells were siRNA-transfected, collected at 48h for DNA fiber analysis. **B.** Immunofluorescence in MCF7 cells 72h after treatment with xentuzumab (100nM) and/or AZD0156 (1μM). Graph: mean ± SEM 53BP1 bodies. **C-D** MCF7 cells treated with 1μM xentuzumab and/or 10μM KU-55933, after 72h processed for: C alkaline come assay; G, neutral comet assay. Left: representative images, right: quantification of % tail DNA (n=100 comets).

As a second approach, we analysed the appearance of 53BP1 bodies in G1 after IGF:ATM co-inhibition, as a readout for a consequence of unresolved replication stress. As previously shown in (Figure 1D), exposure to xentuzumab led to an increase in 53BP1 bodies in G1 (cyclin A-negative) cells. After ATM inhibition there were no detectable 53BP1 bodies, even in combination with IGF-1R inhibition (Figure 8B). This is consistent with reports describing the importance of ATM in phosphorylating the 53BP1 SQ/TQ cluster, required for the subsequent oligomerization of 53BP1 into 53BP1 bodies (29, 57). The absence of 53BP1 bodies after dual ATM:IGF inhibition suggests inadequate protection of under-replicated DNA in replication-stressed IGF-inhibited cells.

Finally, we investigated the nature of the DNA damage induced by IGF-inhibition in the presence and absence of functional ATM, hypothesizing that ATM prevents conversion of tolerable DNA lesions to an intolerable form. If so, this could potentially explain relatively minor inhibitory effects of xentuzumab on viability (Figure 6A) despite significant induction of DNA damage foci in ATM proficient cells (Figure 1A-D). To test this hypothesis, we treated MCF7 cells with xentuzumab, KU-55933 or the combination and quantified SSBs and DSBs by alkaline comet assay, and DSBs only by neutral comet assay. Xentuzumab induced an increase in fragmented DNA in alkaline assays but not neutral assays (Figure 8C-D), suggesting that the damage was predominantly SSBs, comparable to effects of RNR inhibition by hydroxyurea (HU;58). The combination of xentuzumab and KU-55933 significantly increased DNA fragmentation, in excess of the cumulative effects of each drug separately, in both alkaline and neutral assays (Figure 8C-D). This finding suggests appearance of DSBs, potentially originating from SSBs that accumulate after IGF-1R inhibition. Unrepaired DSBs are highly toxic (59), accounting for the significant increase in cell death we had observed in cells exposed to xentuzumab with KU-55933 or AZD0156 (Figure 6G-H). ATM inhibition suppresses 53BP1 body formation, suggesting loss of protection of unreplicated DNA in IGF-inhibited cells, which would account for accumulation of DSBs in IGF:ATM co-inhibited cells, as identified in comet assay (Figure 8C). Our data suggest a new role for ATM in IGF:ATM co-inhibited cells, preventing transition from managed replication stress to toxic DSB accumulation.

## Discussion

RRM2 is the regulatory subunit of the RNR complex, the sole enzyme capable of de novo dNTP synthesis (34). RRM2 expression is reported to be coordinated at the transcriptional, translational and post-translational levels by BRCA1, CDK4/6, mTORC1, WEE1/SETD2 and SCF^Cyclin^ ^F^/CDKs respectively (42, 60-62). We identify IGF-1R as a potent regulator of RRM2 expression, with data to implicate AKT, MEK-ERK and JUN as mediators of the IGF effect on RRM2 expression. Inhibition of IGF-induced RRM2 upregulation in response to AKT inhibitor AZD5363 could be consistent with reported involvement of mTORC1 in RRM2 regulation, although mTORC kinase inhibition was found to downregulate both RRM1 and RRM2 (61), unlike the RRM2-specific effect we observed in IGF-1R depleted cells. Few reports have implicated MEK-ERK in RRM2 regulation: RRM2 was found to be upregulated in KRAS-driven CRC and PDAC cells although the mechanism was not explored, and pharmacologic MEK-ERK inhibition reportedly enhanced radiosensitivity and downregulated DDR proteins including RRM2, also in KRAS-mutant PDAC (20, 63).

RRM2 is frequently overexpressed in cancers and associates with adverse outcomes (64. Therefore, it is of potential therapeutic relevance that its expression and function are regulated by a druggable target. IGFs are well-known to promote cell cycle progression, attributed previously to regulation of G1-S and G2-M checkpoints {Chitnis, 2008 #3). We show here that by regulating RRM2 and replication fork progression, IGFs also influence passage though S-phase. Thus, IGF axis blockade leads to RRM2 downregulation, shortened DNA tracts, RPA and CHK1 phosphorylation and intra-S checkpoint activation. There is a potential discrepancy in our finding of shortened replication tracts and reduced origin firing after IGF-1R depletion, since both of these changes should suppress BrdU incorporation and therefore reduce the S phase fraction. In contrast, we found an increase in the proportion of IGF-1R depleted cells in S phase, and dynamic cell cycle analysis following synchronization showed that Nocodazole trap/release accentuates this S-phase accumulation (Figure 3C-D). The reduced rate of replication we identify, shown by delayed incorporation of BrdU analogs IdU and CIdU (Figure 2A-E), presumably results in delayed transit though S-phase. These findings suggest that the S-phase accumulation we observe upon IGF-1R depletion is explained by longer transit time through S-phase, resolving the apparent discrepancy. IGFs have not been previously linked with regulation of fork progression in the absence of exogenous damage, although replication fork delay induced in murine fibrobasts by alkylating agent methyl methanesulfonate was reportedly exacerbated in IGF-1R null cells, implying that IGF-1R and PCNA are required for DNA damage tolerance (65). The mechanistic basis of delayed fork progression was not discussed in that report, but could relate to our finding that IGF-1 regulates RRM2 expression and dNTP supply. Similarly, IGF-1 was recently reported to rescue from HU-induced fork stalling only in cancer cells where nuclear IGF-1R:PCNA interaction was detected (66).

The replication stress phenotype we identify upon IGF axis blockade contrasts with the well-characterized replication stress associated with positive oncogenic functions. Therefore, this represents the first example of replication stress being induced by inhibition rather than over-activity of an oncogenic RTK. To date, the association of major oncogenic drivers with replication stress has been shown to involve the function of oncogenes including MYC, CYCLIN E and RAS, acting as a positive stimulus to deregulate origin firing (67). Specific molecular mechanisms implicated here include increased origin firing and re-firing leading to re-replication, replication/transcription conflicts that generate R-loops, G-quadruplexes and other secondary structures, or through ROS generation or perturbation of dNTP metabolism (67-71). Ultimately, these initiating factors have been shown to lead to exhaustion of replication substrates including dNTPs (59, 72), suggesting that reduction in the supply of replication substrates may represent a final common pathway that leads to replication stress induced by both oncogene activation and IGF-blockade. Replication stress occurring in the specific context of oncogene-induced senescence has been shown to be due to RRM2 downregulation (73), providing further parallels between the molecular events triggered by oncogene activation and IGF inhibition.

Although the replication stress phenotype induced by IGF blockade was associated with significant slowing of replication fork progression, accumulation of ssDNA lesions and ATR-CHK1 activation, we found only minor reduction in cell viability in xentuzumab treated breast cancer cells. This is reminiscent of the effect of hydroxyurea that targets dNTP synthesis by direct inhibition of RNR, and is reported to cause similarly minor inhibition of cell survival only at relatively high (~100μM) concentration (74). Given evidence of ATR activation in response to under-replicated DNA, we predicted that ATR activation was protective, and ATR inhibition would be toxic to IGF-inhibited cells. However, this proved not to be so (Supplementary Figure S3F-G). Rather, the compound screens highlighted the toxicity of IGF:ATM co-inhibition, leading us to identify a role for ATM in protecting replication forks by preventing conversion of replication stress-associated DNA lesions into DSBs. ATM plays a pyramidal role in regulating the cellular response to DSBs and other cellular stresses by controlling cell cycle checkpoints and coordinating the response to DSB induction (75). DSBs can arise after mishandling of SSBs, and DSB repair through HR or NHEJ is critical because unrepaired DSBs lead to premature arrest of transcription and replication, chromosome breakage and cell death (76). Two novel roles have been described for ATM that are distinct from its classical role as coordinator of DSB response, and which may be relevant here. First, ATM has been shown to be activated in response to unrepaired SSBs, and to be essential for triggering G1 arrest, allowing time for completion of DNA repair to avoid replicating damaged DNA which would result in DSB formation (26). Secondly, ATM is also reportedly activated by RRM2 knockdown, and loss of ATM function was reported to restore dNTP supply in RRM2-depleted cells via metabolic reprogramming to increase substrates for dNTP synthesis, suppressing the hallmarks of replication stress induced by RRM2 depletion, and increasing cellular proliferation (77). However, we found no evidence of ATM activation in the context of RRM2 downregulation and replication stress induced by IGF inhibition, although it is possible that SSBs were induced in IGF-inhibited cells below a threshold reportedly required to activate ATM (26). Furthermore, we found that loss of ATM function in the context of IGF-1R depletion or IGF inhibition was not protective, but rather caused synergistic inhibition of cell viability and enhanced cell death.

We show here that loss of IGF-1R induces a significant delay in replication fork progression, associated with accumulation of ssDNA lesions that are converted to DSBs by co-inhibition of ATM. This transition from managed replication stress to DSB accumulation is new and has not been shown for inhibition of other RTKs. We detect more marked replication fork delay in ATM-deficient CRC cells than in cells expressing functional ATM, suggesting a possible role for ATM in contributing to fork protection in the context of IGF blockade. Many DNA damage proteins have been shown to be required for replication fork protection, including BRCA1/2 and members of the Fanconi Anemia pathway (78-80), although this role has not been previously ascribed to ATM. We propose that the DSBs observed in IGF:ATM co-inhibited cells are due to unprotected unreplicated genomic regions being segregated during the next mitosis, the mechanism being apparently ATR-independent. This newly-identified function of the IGF axis suggests that patients whose tumors lack ATM may be responsive to IGF blockade, and sheds new light on the regulation of global DNA replication.

## Methods

See Supplementary Material for methods including antibodies used in this study (Supplementary Table S1). Study approval: *in vivo* experiments were performed under UK Home Office-approved Project Licence PPL30/3395 and Personal License PILI9BC08CD7.

## Acknowledgements

This study was supported by Breast Cancer Now (2014NovPR364) and Cancer Research UK (C476/A27060), and support to VMM from the NIHR Oxford Biomedical Research Centre and the Harrington Discovery Institute. We acknowledge the assistance and expertise provided by Graham Brown from the Microscopy core facility, Department of Oncology, University of Oxford. We thank Hui Li from Huazhong University of Science and Technology, China for the generous gift of the pGL3-RRM2-firefly construct, Vincenzo D’Angiolella for the kind gift of U2OS cells over-expressing RRM2, Walter F. Bodmer and Jenny Wilding for the gift of HCT15 cells, Sebastian M.B. Nijman for the gift of H322 *ATM* CRISPR cells, Eric O’Neil for the gift of BXPC-3 and Walter F. Bodmer, Amato J. Giaccia and Tim Humphrey for comments on the manuscript.

## Author contribution

GR and VMM designed the study and GR performed most of the experiments, with additional data from XW, LH, JM, AN, LC, ES, SH, DE and LKF. Xentuzumab was provided by UWC and TB, and AJR supervised the design and conduct of *in vivo* experiments. The manuscript was written by GR and VMM and reviewed by all co-authors.

---

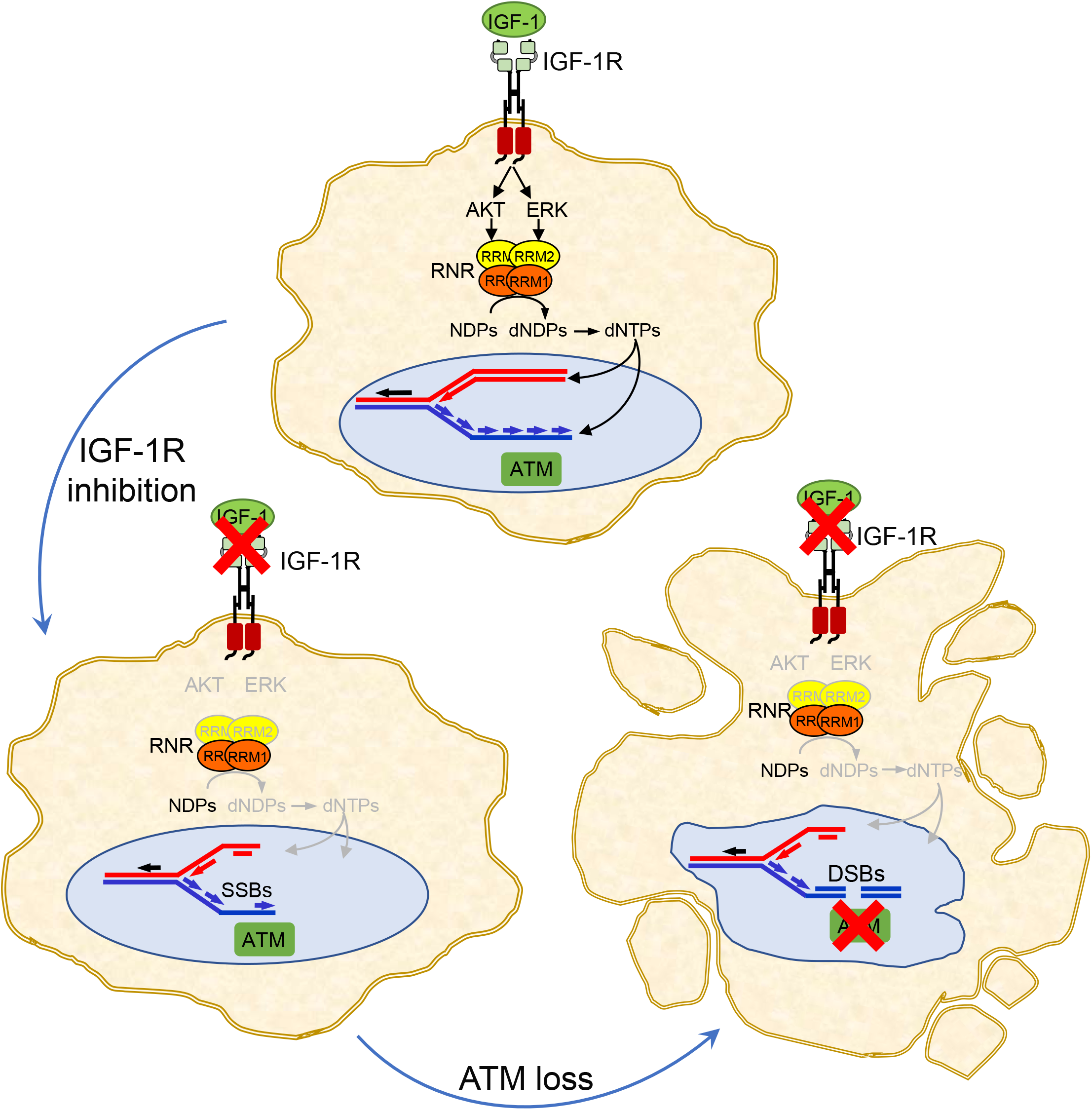

