## Supplementary Figures for "IGF targeting perturbs global replication through ribonucleotide reductase dysfunction"

#### Supp. Figure S1

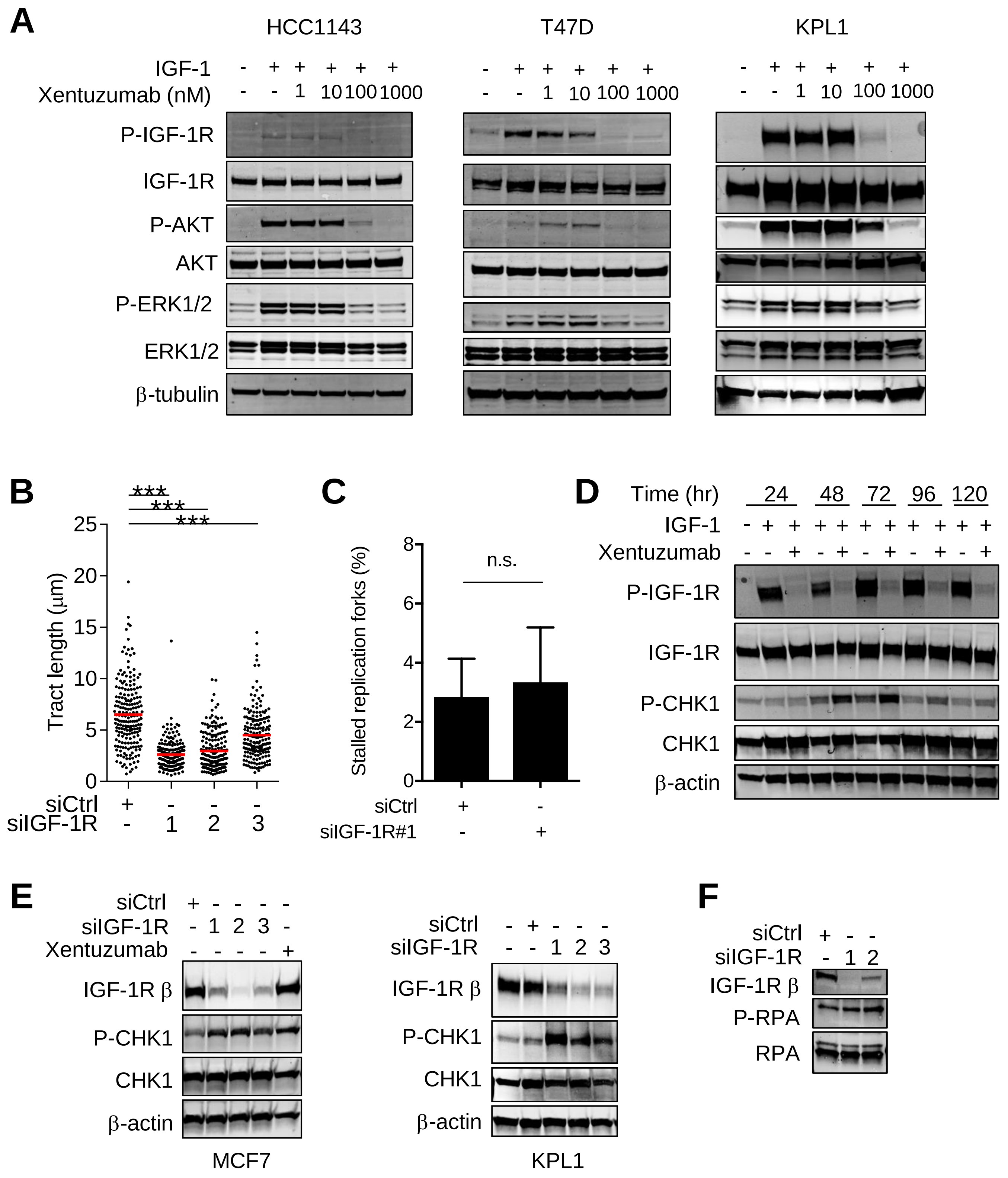

### Supp. Figure S1

**G**

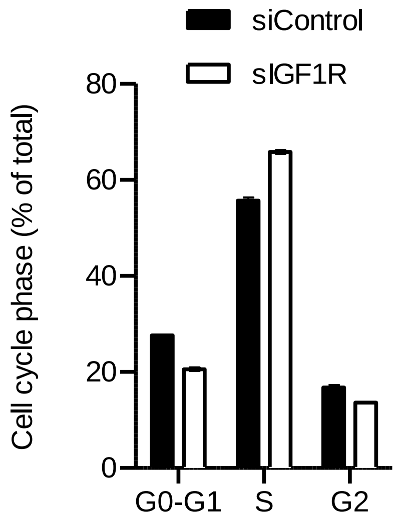


**H**

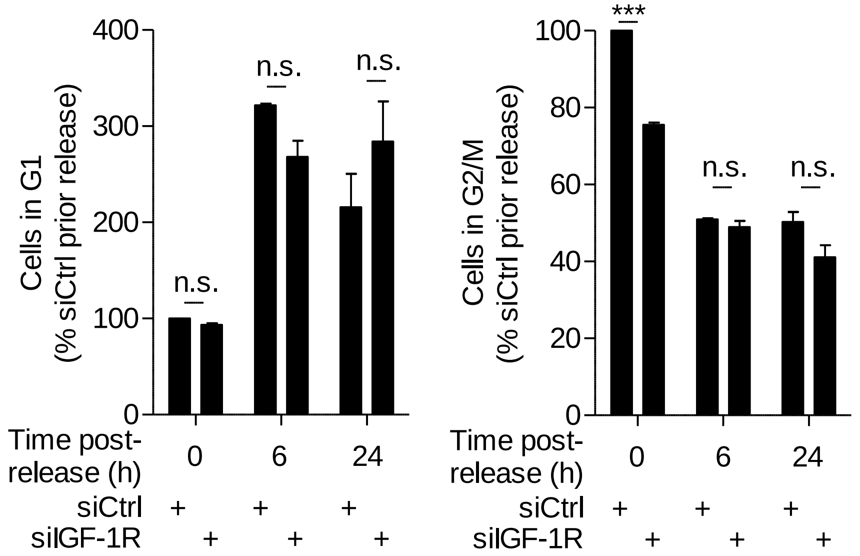


**Supplementary Figure S1. IGF-1R inhibition or depletion slows replication fork progression and activates ATR-dependent CHK1 phosphorylation**. **A.** Serum starved HCC1143, T47D and KPL1 cells were treated with xentuzumab for 5 days and in the final 15 min with 50nM IGF-1. **B.** KPL1 cells transfected with Control (siCtrl) or 3 independent IGF-1R siRNAs were processed 72h later for DNA fiber analysis and replication tract length (CldU/IdU) was quantified. Red line represents the median. **C.** MCF7 cells were siRNA-transfected and analysed by DNA fiber assay as Figure 1E-F, and stalled forks were quantified (n=200 tracts per condition). There was a low percentage of fork stalling, with no difference between siControl and siIGF-1R transfectants. **D.** Serum starved MCF7 cells were treated with xentuzumab for 1-5 days and in the final 15 min with 50nM IGF-1. Cells were lysed for western blot, showing an increase in CHK1 Ser345 phosphorylation in xentuzumab-treated cells, peaking at 72h. **E.** MCF7 (left) and KPL1 cells (right) were transfected with Control (siCtrl) or 3 independent IGF-1R siRNAs and lysed 72h later or treated with 100nM xentuzumab for 72h and lysed for western blot. **F.** MCF7 cells were transfected with Control (siCtrl) or 2 independent IGF-1R siRNAs, and after 72h were lysed for western blot. **G.** MCF7 cells were siRNA-transfected and collected after 48h for cell cycle analysis (n=3 independent experiments). **H.** MCF7 cells siRNA transfected as in Figure 3D were harvested after 42-68h including the nocodazole treatment for cell cycle analysis (n=3 independent experiments).

### Supp. Figure S2

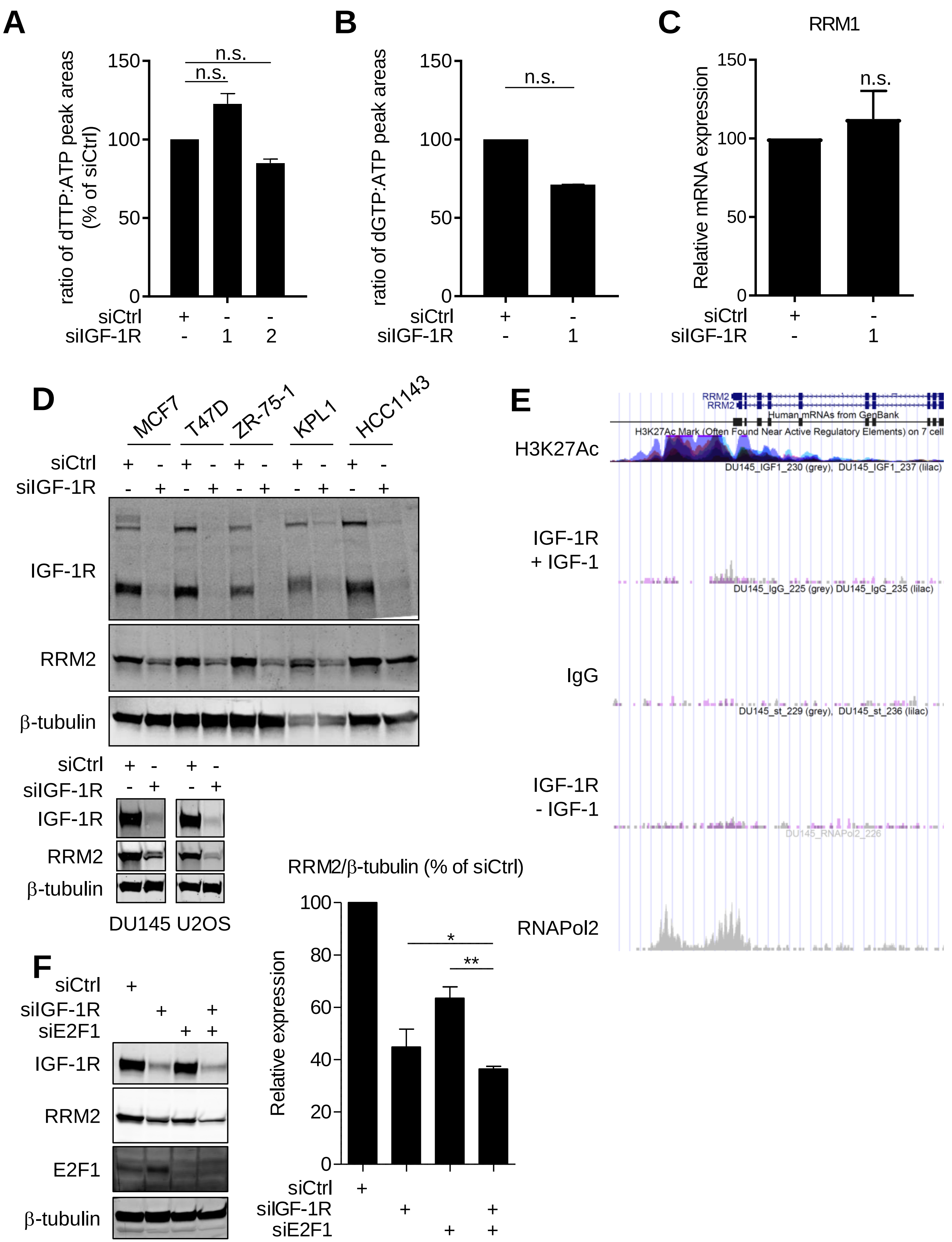

### Supp. Figure S2

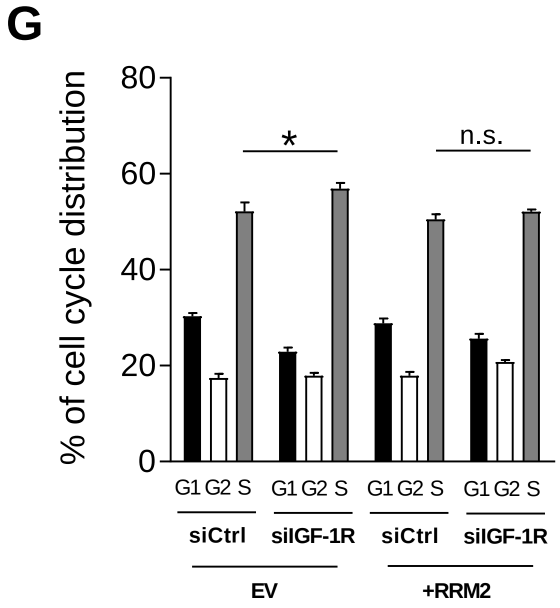


**Supplementary Figure S2. IGF-1R regulates RRM2 levels and RNR activity. A, B.** MCF7 cells were transfected with Control (siCtrl) or IGF-1R siRNAs and after 48h were processed for dTTP (A) and dGTP (B) analysis by HPLC. Mean ± SEM fold change in dTTP and dGTP levels shown as % of siCtrl. **C.** *RRM1* mRNA quantified in MCF7 cells following siRNA-transfection as A-B. **D.** Breast cancer cells MCF7, T47D, ZR-75-1, KPL1, HCC1143, DU145 prostate cancer and U2OS osteosarcoma cells were transfected with Control (siCtrl) or IGF-1R siRNAs and lysed after 48h for western blot to assess IGF-1R depletion and RRM2 expression. **E.** DU145 cells were treated with IGF-1 (IGF-1R + IGF-1) or without (IGF-1R – IGF-1) and analysed by ChIP-seq as we reported [2]. Image from UCSC browser: RRM2 promoter showing enrichment for RNAPol2 recruitment and for H3K27 acetylation, often found near active regulatory elements (public data from ENCODE), but not for IGF-1R (duplicate samples, grey/lilac, or IgG, control ChIP). **F.** MCF7 were siRNA transfected and harvested 48h after transfection for western blot. **G.** MCF7 were stably transfected with a RRM2 vector or empty vector and processed for cell cycle analysis 48h after siRNA transfection.

### Supp. Figure S3

**A**

| Compound | Target |
|---|---|
| KU-55933 | ATM |
| PD318088 | MEK |
| Lapatinib | HER2 |
| BMS-599626 | Pan-HER |
| Fulvestrant | ER |

**B**

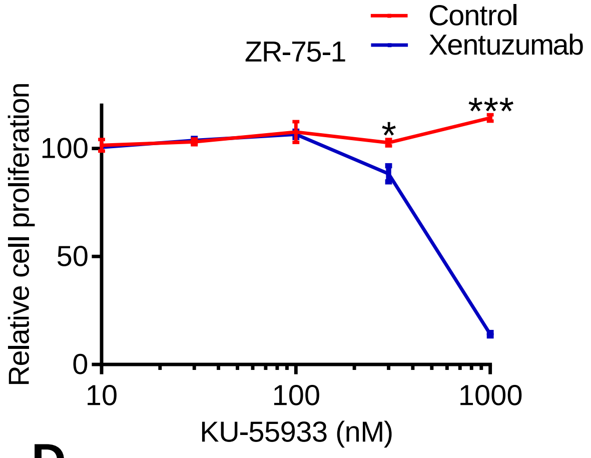


**C**

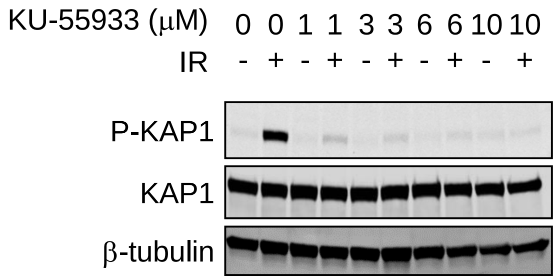


**D**

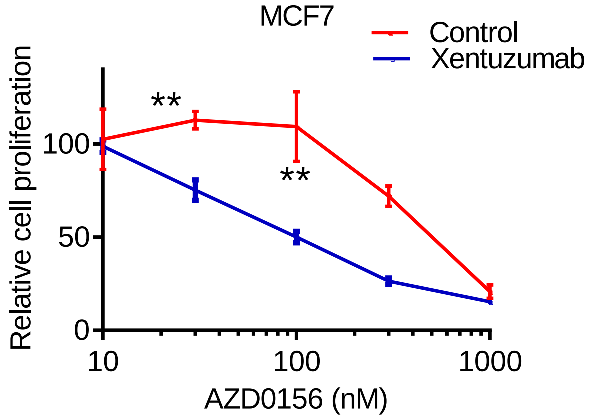


**E**

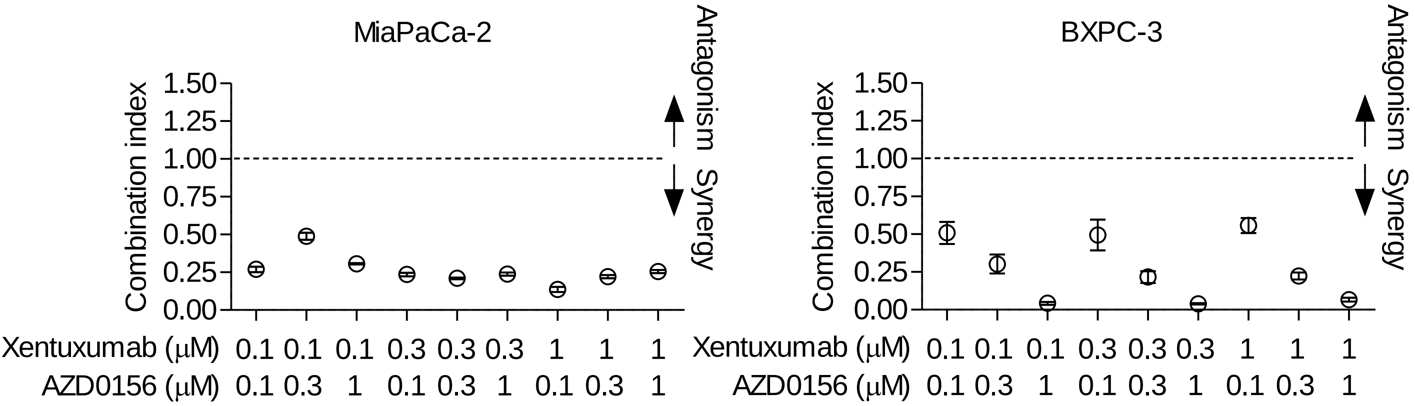


**F**

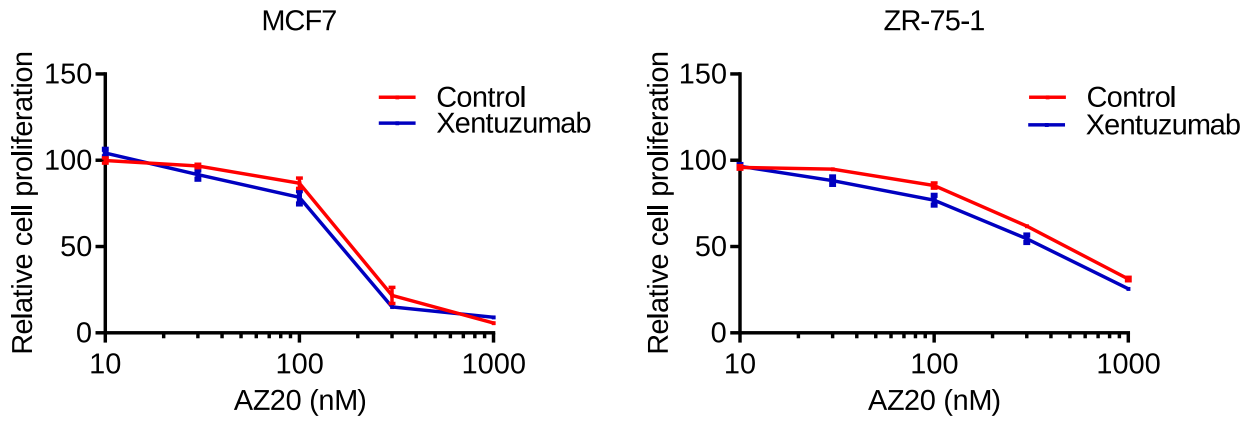

### Supp. Figure S3

G

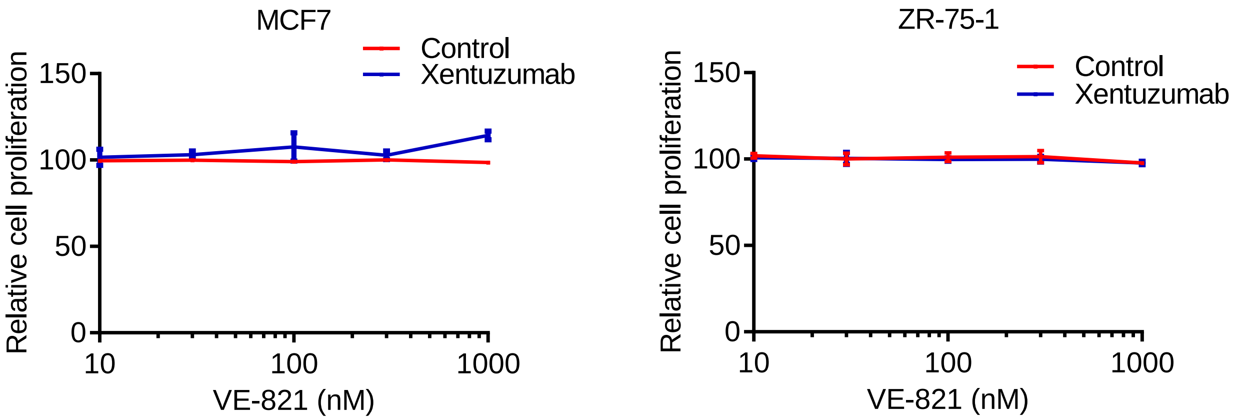


H

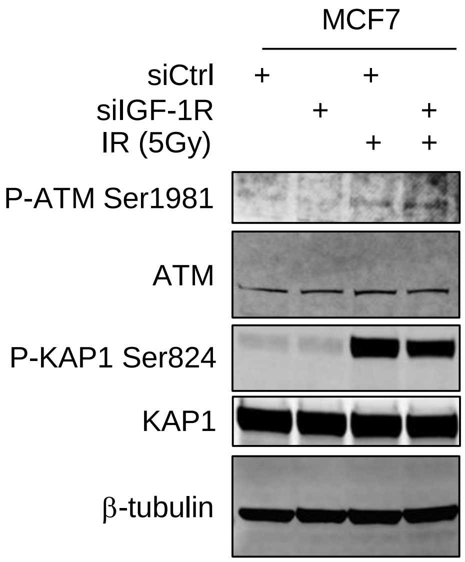


**Supplementary Figure S3. Testing xentuzumab with ATM or ATR inhibitors. A.** Top 5 screen hits identified in common at compound screening concentrations of 0.1, 1 and 10 µM in MCF7 cells. **B.** ZR-75-1 cells were treated with KU-55933 alone or with 1 µM xentuzumab, and viability was measured after 5 days. Xentuzumab induced significant reduction in cell viability. **C.** MCF7 cells were pre-treated with 1-10 µM KU-55933 for 48h, irradiated (5Gy) and collected after 30min for western blot. **D.** MCF7 cells were treated with AZD0156 alone or with 1 µM xentuzumab, and viability was measured after 5 days. **E.** Combination indices (CI) calculated from viability data on MiaPaCa-2 and BXPC-3. CI values <0.8 indicate synergy and <0.3, strong synergy. **F-G.** MCF7 (left) and ZR-75-1 (right) cells treated with ATR inhibitors AZ20 (E) or VE-821 (F) alone or with 1 µM xentuzumab, and viability measured after 5 days. **H.** MCF7 cells pre-transfected with siCtrl or siIGF-1R for 72h, analysed for WB 30 min after 5Gy irradiation.

### Supp. Figure S4

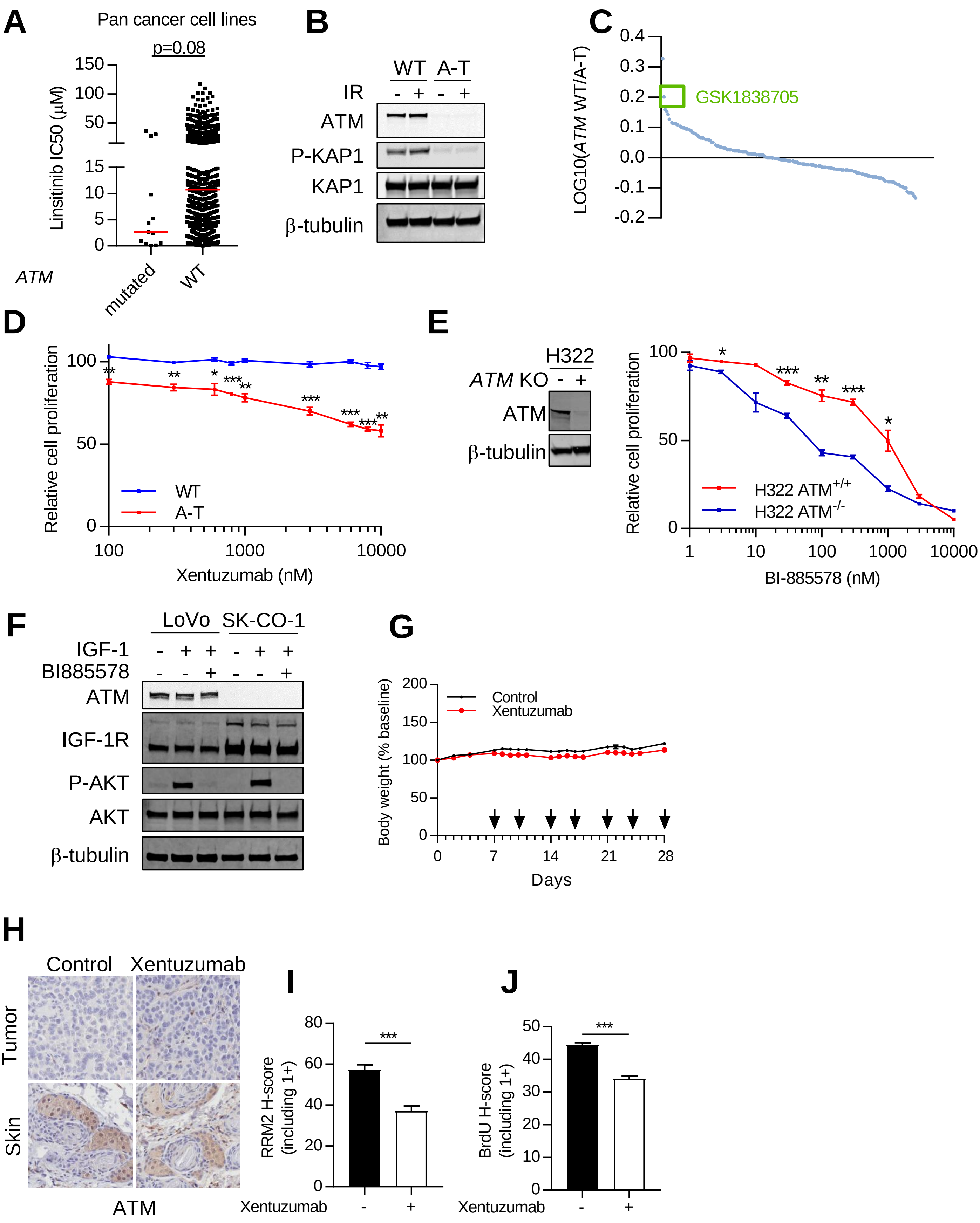

**Supplementary Figure S4. Importance of IGF-1R for *ATM* mutated cells. A.** Pan-cancer analysis on 926 cell lines associating *ATM* homozygous mutations and sensitivity to IGF-1R inhibition with linsitinib (cancerrxgene.org). **B.** *ATM* WT and A-T fibroblasts were irradiated (5Gy) and collected after 30min for western blot. **C.** *ATM* WT and A-T fibroblasts were treated with 0.1 µM kinase inhibitors (188 compounds). Viability was measured by resazurin assay after 3 days. Graph shows results of compounds ranked by ratio of log10 viability *ATM* WT/A-T fibroblasts. Second top hit is highlighted in the green box. **D.** *ATM* WT and A-T fibroblasts were treated with xentuzumab and viability was measured after 5 days. **E.** ATM level in H322 isogenic CRISPR cells. **F.** Serum starved LoVo and SK-CO-1 cells were pre-treated with BI-885578 for 24h and in the final 15 min with 50nM IGF-1. **G.** Body weight of mice from figure 4F bearing SK-CO-1 xenografts treated twice weekly (arrows) with solvent or xentuzumab. **H.** Immunohistochemical staining for ATM of SK-CO-1 tumors treated with control or xentuzumab for 3 weeks. **I-J.** Quantification of immunohistochemical signal for RRM2 (H) and BrdU (I) from figure 7G, including weak positive (1+) cells.
