## Supplementary Tables and Methods for "IGF targeting perturbs global replication through ribonucleotide reductase dysfunction"

**Supplementary methods**

**Cell lines and reagents.** Breast cancer cell lines HCC1143, MCF7, T47D and ZR-75-1 were obtained from Anthony Kong (Institute of Cancer and Genomic Sciences, University of Birmingham), HCT15 cells from Walter Bodmer (Department of Oncology, University of Oxford), BXPC-3 from Eric O'Neil (Department of Oncology, University of Oxford), *ATM* WT and *ATM* null H322 cells from Sebastian Nijman, Nuffield Department of Medicine, University of Oxford and KPL1, LoVo, SK-CO-1, immortalized *ATM* WT MRC5-SV2 from the European Collection of Authenticated Cell Cultures (ECACC) and $ATM^{-/-}$ AT5BIVA fibroblasts from The Coriell Institute for Medical Research, MiaPaCa-2 from the American Type Culture Collection (ATCC). Cell lines were mycoplasma-free when tested with MycoAlert (Lonza Rockland Inc.). Cell line identity was authenticated by STR genotyping (Eurofins Medigenomix Forensik GmbH). Cells were grown in DMEM (Thermo Fisher Scientific) supplemented with 10% fetal calf serum (FCS) (Invitrogen), penicillin (100 units/mL) and streptomycin (0.1mg/mL; Invitrogen) at 37°C in a humidified incubator with 5% $CO_2$.

Xentuzumab and BI-885578 were from Boehringer Ingelheim, other inhibitors as follows: KU-55933 (Bio-Techne; R&D systems), trametinib and AZD0156 (Selleck), AZD5363 and 60 drug custom library (Stratech Scientific), 188 compound kinase chemogenomic set (KCGS, Structural Genomics Consortium; www.sgc-unc.org/kinase-chemogenomics). Cells were transfected with 50 nM IGF-1R siRNAs from Qiagen, Cell Signaling Technology or generated in-house (81) or with AllStars Negative Control siRNA (Qiagen), using Oligofectamine (Invitrogen) as described in (39).

**Cell viability and death assays.** Cells seeded in 96 well plates were treated with drug(s) and after 5 days underwent viability assay (CellTiter-Blue, Promega) measuring fluorescence (570nm) on a POLARstar Omega microplate reader (BMG Labtech). Viability data were used to calculate Combination Indices using ComboSyn software following the Chou-Talalay method (82). For death assay, cells were seeded and treated as above and stained with 1mg/ml Hoechst 33342 and 4 µM propidium iodide (PI) to quantify total and dead cells on a Celigo Imaging Cytometer (Nexcelom).

**Western blotting** was performed as previously (83). Bound primary antibodies (Supplementary Table S1) were detected by secondary antibodies coupled to IRDye (LI-COR), visualised using the Odyssey Quantitative Fluorescence Imaging System, and band intensities were quantified using Odyssey software.

**Immunofluorescence**. Cells were seeded and treated in 4-well Labtek plates and were fixed for 10 min in 4% paraformaldehyde (PFA), washed in PBS and permeabilized for 5 min in PBS 0.1% Triton/4% FCS. Cells were washed in PBS, blocked for 1h in 10% BSA, washed in PBS and incubated with primary antibodies (Supplementary Table S1) for 1h at RT. Cells were washed in PBS, incubated with secondary antibodies for 1h at RT in the dark. Slides were air-dried and mounted in VectaShield (H1200, Vector Laboratories) with DAPI. Images acquisitions were performed using an inverted laser scanning confocal LSM 710 microscope UV Zeiss equipped with x63 and x40 oil objectives. Images were processed with the Zeiss software (Zen), imported to ImageJ, pseudocolored and merged.

**DNA fiber assays** were performed as described (84, 85). Briefly, cultures in exponential growth were incubated with CldU (25 µM, Sigma, C6891) for 20 min. Cells were washed 3 times in warm PBS and incubated in medium containing IdU (250 µM, Sigma, I7125) for 20 min. Cells were lysed and nuclei were spread onto glass slides in spreading buffer (200 mM Tris-HCl pH 7.4, 50 mM EDTA, 0.5% SDS). Slides were slightly tilted to allow DNA to spread slowly on the whole length of the slide, and then air dried and fixed with methanol/acetic acid (3:1) for 10min. To detect incorporation of CldU/IdU, slides were washed in PBS, DNA was denatured in 2.5 M HCl (1 h, RT) and blocked in 2% BSA+ 0.1% Tween and incubated (overnight, 4°C) with mouse anti-IdU antibody and rat anti-CIdU in blocking solution. Slides were washed 5 times in PBS+Tween 0.2%, briefly rinsed in blocking solution and incubated (1 h, RT) with anti-mouse Alexa Fluor 488 and anti-rat Alexa Fluor 594 antibodies in blocking solution. Slides were washed 5 times in PBS+Tween 0.2% and mounted in Vectashield. Fork velocity was converted to kb/min based on the tract length (µm) and the duration of CIdU/IdU incubation (20min each).

**Flow cytometry**: Following siRNA transfection and nocodazole treatment (Sigma-Aldrich M1404, 60ng/mL), cultures in exponential growth were pulsed with BrdU (20µM, Invitrogen) for 30 min. Monolayer and suspension cells were collected and processed as in (16, 21)**.** In brief, cells were fixed in 70% ethanol for 30 min, denatured in 2M HCl containing 0.1 mg/mL pepsin for 20 min. After washes with PBS and 2% FCS, cells were incubated with primary anti-BrdU antibody (1:500, BD Bioscience, 347580) overnight at 4°C. After washes with PBS and 2% FCS, cells were incubated with secondary antibody anti-mouse Alexa Fluor 488 (1/500, Molecular Probes) for 1h at RT. After PBS washes, cells were resuspended in 0.1 mg/ml PI. Fluorescence was analysed using a FACSCalibur instrument (BD Biosciences), and results were analyzed with FlowJo software (v.10).

**Nucleotide quantification.** Cells were harvested and nucleotides extracted with 70% ice-cold methanol. Precipitated proteins were removed by centrifugation, supernatants were stored at −80 °C then dried in a heated vacuum centrifuge and reconstituted in HPLC starting eluent. Determination of the dNTP pool size was performed by HPLC as described in (71).

**Quantitative PCR**: RNAs were extracted and reverse transcribed using the Pure Link RNA Mini RNA extraction kit (Ambion) and SuperScript III First-Strand Synthesis SuperMix (Invitrogen). The cDNAs were amplified using primers RRM1 F (AAACGCCATCGCCCCATTGGAA), RRM1 R (ACAGCTGGCTTCCAGAGCACCATA), RRM2 F (TTTAAAGGCTGCTGGAGTGAGG), RRM2 R (GCAGCTGCTTTAGTTTTCGGCT), β-actin F (AGGATTCCTATGTGGGCGAC) and β-actin R (ATAGCACAGCCTGGATAGCAA) and Sybr Green PCR Mix (Applied Biosystems) on a 7500 Fast RT-PCR System (Applied Biosystems).

**Luciferase Assays:** ONE-Glo™ EX Luciferase assay (Promega Cat. No. E8110) was used for testing of the effects of IGF-1R, c-JUN and E2F1 knockdown on RRM2 transcription levels. Briefly, MCF7 cells were seeded to be 70% confluent for transfection, then transfected with the RRM2 promoter reporter construct using Lipofectamine 3000™ (Invitrogen Cat. No. L3000008) as per the manufacturer's instructions. Control wells were also transfected with only a constitutively expressed GFP plasmid as a transfection efficiency control. SiRNAs for IGF-1R, c-Jun and E2F1 or All-Stars control were transfected 24h later and, following overnight incubation at 37°C, cells were reseeded in triplicate into a 96-well plate. The following day cells were tested with ONE-Glo™ EX Luciferase assay reagent (Promega Cat. No. E8110). Briefly, treated cells and ONE-Glo™ EX Luciferase assay reagent were equilibrated to RT before addition of ONE-Glo™ EX Luciferase assay reagent in 1:1 ratio of reagent to medium. Cells were incubated in luciferase reagent for 5 mins, RT on a rocker and shielded from light, and luminescence was measured on a POLARstar Omega plate reader (BMG Labtech). The protein content of the lysate was then quantified by Pierce BCA protein assay (ThermoFisher Cat. No. 23225) and luciferase activity was expressed as relative light units per µg protein.

**Generation of MCF7 stably overexpressing RRM2:** To generate RRM2-overexpressing MCF7 cells, pcDNA 3.1 control plasmid (#V79520, Invitogen) or pcDNA3.1 RRM2 plasmid (Addgene, Plasmid

#13796) was transfected into MCF7 cells using Lipofectamine 3000 Reagent Kit (#L3000001, Thermo Fisher Scientific) following the manufacturer's instructions. MCF7 cells were cultured to 70% confluency. Lipofectamine 3000 reagent and P3000 reagent were diluted with Opi-MEM medium (#31985-047, Gibco) and then mixed with 2.5 µg plasmid DNA. The transfection mix was added into cell culture medium for 48 hours. The cells were then cultured with medium containing 800 µg/mL G418 (#G8186, Sigma-Aldrich) for 72 hours. The surviving colonies were picked and cultured with medium containing 800 µg/mL G418 for 30 days to obtain stably expressing cells. The expression level of RRM2 was determined by western blot analysis.

**Compound screens**: Two compound screens were performed, first to test xentuzumab with a targeted custom library in breast cancer cells, and secondly to test a library of kinase inhibitors in *ATM* WT MRC5-SV2 vs *ATM*$^{-/-}$ AT5BIVA fibroblasts. Duplicate screens were conducted in each case. For the breast cancer screen, 60 compounds were tested at 100 nM, 1 µM and 10 µM alone or in combination with 1 µM xentuzumab in MCF7, HCC1143, KPL1, T47D and ZR-75-1 cells. The cells were seeded at pre-optimised cell densities in 96-well plates by Perkin Elmer Janus liquid handling workstations. Cells were incubated for 24h at 37°C and treated with a drug library of 60 cell cycle, replication and DNA repair inhibitors alone or with 1 µM xentuzumab using the Perkin Elmer Janus liquid handling workstations. Cells were incubated for 5 days at 37°C followed by replacement of the medium by phenol-free DMEM containing 1mM resazurin (Cell Titer Blue, Promega). Cells were incubated for 2h at 37°C and fluorescence was read using EnVision multilabel plate reader (PerkinElmer). Screen replicate correlation was 0.97 and the screen Z-Factor calculated using the positive control for viability inhibition (PLK1 inhibitor BI-2536) was 0.64 at 100 nM, 0.68 at 1 µM and 0.72 at 10 µM, indicating a high-quality screen. The compound screen in *ATM* WT vs null fibroblasts used a 188-compound kinase chemogenomic set (KCGS), and viability was measured after 3 days as above. The screen Z-Factor calculated using the positive control for viability inhibition (camptothecin analogue SN-38) was 0.63 and 0.81 at 0.1 µM for *ATM* WT and mutated cells respectively, and 0.83 and 0.81 at 1 µM, indicating a high-quality screen.

**Single cell gel electrophoresis.** Alkaline comet assays were performed as in (86, 87). Briefly trypsinised cells were mixed with 1% low melting point agarose, spread onto glass slides on ice and flattened with cover slips. After solidification, cover slips were removed and slides were lysed in lysis buffer (2.5M NaCl, 100mM EDTA, 10mM Tris pH10.5, with *ex tempo* addition of 1% Triton X-100 and 1% DMSO) for 1h at 4°C. Slides were placed in an electrophoresis tank with fresh electrophoresis buffer (300mM NaOH, 1mM EDTA and 1% DMSO) for 40min to allow DNA unwinding. Slides were then subjected to electrophoresis for 25min at 25V, 300mA at RT. Slides were rinsed with neutralisation buffer (TrisHCl 500mM pH 8), stained with SyBr Gold (ThermoFisher) and stored at 4°C before image acquisition. For neutral comet assays, cells were agarose-embedded and spread on slides as above, and lysed in lysis buffer (2.5M NaCl, 100mM EDTA, 10mM Tris and 1% N-lauroylsarcosine pH9.5, with *ex tempo* addition of 1% Triton X-100 and 1% DMSO) for 2h at 4°C. Slides were placed in an electrophoresis tank, incubated with fresh TBE for 30min to allow DNA to unwind, and subjected to electrophoresis for 32min at 72V at RT. Slides were rinsed, stained and stored as above. Acquisition and analysis of 100 comets per condition were performed at 100× magnification using an epifluorescence microscope (Ni-E, Nikon) and Komet v.5.5 software (Andor). DNA damage was measured by quantifying tail DNA as % total DNA.

***In vivo* study.** Experiments were performed under UK Home Office approved project licence (PPL) 30/3395. SK-CO-1 cells ($10^7$/mouse in 50% Matrigel, BD Bioscience) were injected into the subcutaneous tissues of the back of 6 week-old female CD-1 nude mice (Envigo). When tumors reached 100 $mm^3$, mice were randomly divided into two groups (n=10) for treatment with control (PBS) or 100 mg/kg xentuzumab twice weekly by intraperitoneal injection. Body weights and tumor volumes were measured 5 times a week for 3 weeks. On the final day, mice were injected intraperitoneally with BrdU, and after 2h mice were humanely culled and tissues processed for IHC.

**Immunohistochemistry.** Murine tissues were used under PPL 30/3395 and PIL I9BC08CD7. Freshly cut 6 µm sections of formalin-fixed paraffin embedded tumor and overlying mouse skin were stained using EnVision G2 Double stain System (Agilent Technologies) according to manufacturers' instructions, with additional blocking using 3% $H_2O_2$ for 20min and mouse-on-mouse blocking for mouse antibodies (MKB-2213, Vector Laboratories), and with primary antibodies listed in Supplementary Table S1.

**Statistics.** GraphPad Prism v7 was used to perform Student's t-test to compare means of 2 groups, and one-way ANOVA with Tukey post-test to compare >2 groups. All tests were 2-sided and p<0.05 (*), p<0.01 (**) and p<0.001 (***) were considered significant.

| Western blotting and immunofluorescence | | | | |
|---|---|---|---|---|
| **Primary antibodies** | **Supplier** | **Species** | **Dilution** | **Catalogue number** |
| P-IGF-1R Tyr1135/1136 | Cell Signaling Technology | Rabbit | 1/1000 | #3024 |
| IGF-1R β | Cell Signaling Technology | Rabbit | 1/1000 | #3027 |
| P-AKT Ser473 | Cell Signaling Technology | Rabbit | 1/1000 | #4060 |
| AKT | Cell Signaling Technology | Rabbit | 1/1000 | #9272 |
| P-CHK1 Ser345 | Cell Signaling Technology | Rabbit | 1/1000 | #2348 |
| CHK1 | Santa Cruz | Mouse | 1/1000 | sc-8408 |
| RRM1 | Santa Cruz | Goat | 1/1000 | sc-11733 |
| RRM2 | Santa Cruz | Rabbit | 1/1000 | sc-10844 |
| P-ERK Thr202/Tyr204 | Cell Signaling Technology | Rabbit | 1/1000 | #9101 |
| ERK | Cell Signaling Technology | Mouse | 1/1000 | #4696 |
| P-RPA Ser33 | Bethyl Laboratories | Rabbit | 1/1000 | A300-246A |
| RPA | Cell Signaling Technology | Goat | 1/1000 | #2208 |
| P-S6 Ser235/236 | Cell Signaling Technology | Rabbit | 1/1000 | #2211 |
| S6 | Cell Signaling Technology | Rabbit | 1/1000 | #2217 |
| P-KAP1 Ser824 | Bethyl Laboratories | Rabbit | 1/1000 | A300-767A |
| KAP1 | Bethyl Laboratories | Rabbit | 1/1000 | A300-275A |
| β-actin | Sigma | Mouse | 1/5000 | A2228 |
| β-tubulin | Sigma | Mouse | 1/5000 | T4026 |
| gH2AX | Millipore | Mouse | 1/500 | 05-636 |
| 53BP1 | Novus Biologicals | Rabbit | 1/500 | NB-100-304 |
| Cyclin A | Santa Cruz | Mouse | 1/100 | sc-271682 |
| IdU | BD Bioscience | Mouse | 1/100 | 347580 |
| CIdU | Abcam | Rat | 1/500 | ab6326 |
| **Secondary antibodies** | **Supplier** | **Species** | **Dilution** | **Catalogue number** |
| IRDye 680CW anti-mouse | LI-COR | Mouse | 1/15000 | 926-32212 |
| IRDye 800CW anti-rabbit | LI-COR | Rabbit | 1/10000 | 926-32213 |

| | | | | |
|---|---|---|---|---|
| AlexaFluor 488 | ThermoFisher | Mouse | 1/2000 | A-11001 |
| AlexaFluor 594 | ThermoFisher | Rabbit | 1/2000 | R37117 |
| AlexaFluor 594 | ThermoFisher | Rat | 1/300 | A-11007 |
| **Immunohistochemistry** | | | | |
| **Primary antibodies** | **Supplier** | **Species** | **Dilution** | **Catalogue number** |
| P-AKT Ser473 | Cell Signaling Technology | Rabbit | 1/100 | #4060 |
| Cleaved Caspase 3 | Cell Signaling Technology | Rabbit | 1/150 | #9661 |
| RRM2 | Santa Cruz | Goat | 1/400 | sc-10846 |
| BrdU | BD Bioscience | Mouse | 1/50 | 347580 |
| ATM | Abcam | Rabbit | 1/300 | ab3280 |

**Supplementary Table S1.** Antibodies used in western blotting, immunofluorescence and IHC.

**A. 0.1 µM**

| Rank | Target | Compound |
|---|---|---|
| 1 | AKT | MK-2206 |
| 2 | EGFR | Lapatinib Ditosylate |
| 3 | CDK4/6 | Abemaciclib (LY2835219) |
| 4 | HADC | CUDC-101 |
| 5 | EGFR | BMS-599626 |
| 6 | MEK | PD318088 |
| 7 | Topoisomerase I | Irinotecan |
| 8 | ATM | KU-55933 |
| 9 | Estrogen receptor | Fulvestrant |
| 10 | MDM2 | Nutlin 3 |
| 11 | PI3K, DNA-PK | PIK-75 |
| 12 | DNA/RNA synthesis | Flucytosine |

**B. 1 µM**

| Rank | Target | Compound |
|---|---|---|
| 1 | EGFR | Lapatinib Ditosylate |
| 2 | EGFR | CUDC-101 |
| 3 | CDK1 | Ro-3306 |
| 4 | EGFR | BMS-599626 |
| 5 | RAD51 | BO2 |
| 6 | MEK | PD318088 |
| 7 | ATM | KU-55933 |
| 8 | Mdm2/p53 | Nutlin 3 |
| 9 | Estrogen receptor | Fulvestrant |
| 10 | Aurora kinase | Tozasertib (VX-680) |
| 11 | ATM | KU-60019 |

| | | |
|---|---|---|
| 12 | CDK | PD0332991 |

**C. 10 µM**

| Rank | Target | Drug name |
|---|---|---|
| 1 | ATM | KU-55933 |
| 2 | MEK | PD318088 |
| 3 | EGFR | Lapatinib Ditosylate |
| 4 | EGFR | BMS-599626 |
| 5 | Estrogen receptor | Fulvestrant |
| 6 | MRE11 | Mirin |
| 7 | MEK | BIX 02189 |
| 8 | PARP | ABT-888 |
| 9 | DNA/RNA synthesis | Flucytosine (Ancobon) |
| 10 | mTOR | Deforolimus (MK-8669) |
| 11 | WRN | NSC 19630 |
| 12 | Aurora kinase | Hesperadin |

**Supplementary Table S2. Compound screen in MCF7 cells with xentuzumab.** Tables show top ranked screen hits in cells treated with 1 µM xentuzumab plus compounds at 0.1, 1.0 or 10 µM.
